# Evaluation of the minimum sampling design for population genomic and microsatellite studies. An analysis based on wild maize

**DOI:** 10.1101/2020.03.06.980888

**Authors:** Jonás A. Aguirre-Liguori, Javier A. Luna-Sánchez, Jaime Gasca-Pineda, Luis E. Eguiarte

## Abstract

Massive parallel sequencing is revolutionizing the field of molecular ecology by allowing to understand better the evolutionary history of populations and species, and to detect genomic regions that could be under selection. However, the needed economic and computational resources generate a tradeoff between the amount of loci that can be obtained and the number of populations or individuals that can be sequenced. In this work, we analyzed and compared two extensive genomic and one large microsatellite datasets consisting of empirical data. We generated different subsampling designs by changing the number of loci, individuals, populations and individuals per population to test for deviations in classic population genetics parameters (*H*_S_, *F*_IS_, *F*_ST_) and landscape genetic tests (isolation by distance and environment, central abundance hypothesis). We also tested the effect of sampling different number of populations in the detection of outlier SNPs. We found that the microsatellite dataset is very sensitive to the number of individuals sampled when obtaining summary statistics. *F_IS_* was particularly sensitive to a low sampling of individuals in the genomic and microsatellite datasets. For the genomic datasets, we found that as long as many populations are sampled, few individuals and loci are needed. For all datasets we found that increasing the number of population sampled is important to obtain precise landscape genetic estimates. Finally, we corroborated that outlier tests are sensitive to the number of populations sampled. We conclude by proposing different sampling designs depending on the objectives.

## INTRODUCTION

Massive parallel sequencing (MPS) has revolutionized the fields of molecular ecology, population genetics and landscape genetics (Stapley et al., 2010; Ekblom and Galindo, 2010; Scholz et al., 2012). By increasing the number of polymorphic sites, it is now possible to estimate with higher resolution the genetic diversity, genetic structure and demographic history of populations (Allendorf et al., 2010, Schoville et al., 2012; Excoffier et al., 2013; Meier et al., 2017; Aguirre-Liguori et al., 2019a), and the environmental and geographic mechanisms that determine the connectivity between populations (Bradburg et al., 2013). MPS also allows identifying genomic regions that could be under selection (Foll & Gaggiotti; 2008; Coop et al., 2012; Stapley et al., 2012; De Villemereuil & Gaggiotti, 2015).

MPS is powerful to detect patterns of local adaptation and understanding how environment structures genetic diversity; nevertheless, its potential capacity depends on sampling a large geographic area, encompassing an adequate environmental and genomic representation of the species (Schoville et al., 2012; de Mita et al., 2013; Tiffin & Ross-Ibarra 2014). Unfortunately, for many research groups MPS is still expensive, or in some other cases -such as rare or endangered species-, obtaining large number of populations or individuals, and/or enough DNA for genomic studies can be challenging. In addition, the bio-informatic processing required for large samples can be limiting, making it difficult to obtain adequate genomic representation for enough individuals and populations. A solution has been to prioritize sequencing power to compensate for fewer individuals or populations. However in the context of local adaptation sampling populations growing in different parts of the distribution or different environments can affect the adequate estimation of genetic parameters (Meirmans, 2015), for instance if edge populations are not sampled, and the detection of outlier regions that might be under selection (de Mita et al., 2013). Thus it is crucial to determine the potential biases associated to sampling effort (number of individuals, loci and populations) and to define the trade-off between sampling effort and number of polymorphic regions obtained with MPS to reduce such biases and to obtain robust estimates (Pruett & Winker, 2008; Fumagalli, 2013; Willing et al., 2012; de Mita et al., 2013).

So far, different studies have evaluated the errors and biases generated in estimates of genetic parameters when different number of populations, number of polymorphic sites and number of individuals are used (Table 1 summarizes a list of studies that have evaluated sampling designs on population genetics studies). In summary, these studies have shown that parameters of mean genetic diversity (*F_ST_, F_IS_, H*s) are not significantly affected by sampling different number of loci, number of individuals or number of populations; however, the variance decreases as the number of populations, individuals and loci increases (References in Table 1). In contrast, these studies (References in Table 1) have shown that patterns of isolation by distance and isolation by environment across reduced areas are sensitive to the number of populations sampled and sampling design (linear, aggregated, random sampling).

**TABLE 1.** Summary of 19 studies that have evaluated sampling designs using different markers (Microsatellites, AFLPs, SNPs); empirical vs. simulated data; and varying the number of loci, individuals and populations.

| Reference | Dataset | Type of sampling | N° of populations | N° of individuals | N° of Loci | Principal conclusions |
| --- | --- | --- | --- | --- | --- | --- |
| Miyamoto et al., 2008 | Microsatellite | Empirical | 1 | 480 | 4 | >30 individuals increases the precision in $H_s$<br>Between 200 and 300 individuals increase the precision of allelic richness estimates |
| Pruett & Winker 2008 | Microsatellite | Empirical | 1 | 200 | 8 | Precision in summary statistics is increased when > 20 individuals are genotyped |
| González-Ramos et al., 2015 | Microsatellite | Empirical | 2 | 64 | 15 | Above 6 polymorphic markers are enough to define adequately the genetic structure between populations |
| Peterman et al., 2016 | Microsatellite | Empirical | 5 | 80 | 15 | Increasing the number of loci does not change mean summary statistics, but increases the precision across replicates. IBD patterns are sensitive to few loci genotyped. |
| Sánchez-Montes et al., 2017 | Microsatellite | Empirical | 17-21 (different species) | 547, 652 y 516 | 18, 16 y 15 | > 20 individuals and between 50 and 80 individuals per population are needed to estimate with precision $H_s$, and allelic richness, respectively. |
| Rico, 2017 | Microsatellite | Simulation | 17 and 34 (different species) | 5,000 y 3,000 | 20 | Spatial sampling design (random, systemic, cluster) affect IBD patterns. Increasing loci, over individuals, increases the accuracy of IBD estimates. |
| Schwartz & McKelvey, 2009 | Microsatellite | Simulation | 1 | 10,000 | 15 | Different sampling designs generate different $F_{ST}$ estimates, and different *Structure* outputs. |
| Landguth et al., 2012 | Microsatellite | Simulation | 1 | 1000 | 25 | Increasing the number of polymorphic loci increases the precision of patterns of isolation by resistance (IBR) |
| Oyler-McCance et al., 2013 | Microsatellite | Simulation | 1 | 1000 | 25 | Increasing the number of polymorphic loci, individuals and number of alleles increases the precision and the accurate estimation of patterns of (IBR) |
| Landguth y Schwartz 2014 | Microsatellite | Simulation | 64 | 64 | 20 | Increasing the number of populations (even if fewer individuals are sampled) increases the possibility of finding correct patterns of IBD. |
| Smith & Wang, 2014 | Microsatellite | Simulation | 3 | 100 | 100 | Reducing the number of samples do not affect $H_s$, $F_{ST}$ estimates, but reduces the power to detect accurate allelic richness. |
| Hale et al., 2012 | Microsatellite | Mixed (Simulation and empirical) | 4 | 100 | 9, 5, 7 y 8 | For four different species, sampling between 25-30 individuals are enough to estimate accurately $H_S$ and $F_{ST}$ |
| Dubois *et al.*, 2017 | Microsatellite | Mixed (Simulation and empirical) | 4 | 4 different taxa: 726, 408, 372, 384 | 16 | Sex proportions do not affect summary statistics estimates. > 20 individuals increase the precision of summary statistics. Empirical and simulated data show different patterns of deviation. |
| Sinclair & Hobbs, 2009 | AFLPs | Empirical | 6 | 159 | 59 y 117 | > 30 individuals per population needed to estimate accurately $F_{ST}$ |
| Willing et al., 2012 | SNPs | Simulation | 2 | 1000 | 21,000 | Fewer individuals are needed to estimate accurately $F_{ST}$ for MPS datasets |
| Fumagalli, 2013 | SNPs | Simulation | 1 | 1000 | 20,000 | Low individual sampling, with a high genome coverage underestimates the number of segregating sites, $H_S$ estimates and genetic structure |
| Nazareno et al., 2017 | SNPs | Empirical | 2 | 70 | 3,500 | Few individuals (8) but with a large number of SNPs (> 1000) increase the precision of $H_S$ and $F_{ST}$. |
| Flesch *et al.*, 2018 | SNPs | Empirical | 4 | 120 | 14,000 | > 25 individuals (with 10,000 SNPs) are needed to estimate accurate kinship indexes (10, 000 SNPs), identify identical by descent alleles and $F_{ST}$ values. |
| Puckett y Eggert, 2016 | Mixed (SNPs and Microsatellite) | Empirical | 34 | Microsatellites dataset: 506<br>SNP dataset: 96 | Microsatellite dataset: 15<br>SNP dataset: 1,000 | 1000 SNPs are more precise than microsatellites for assigning birth areas, even if few individuals are sampled. |

Sampling design has been studied widely. Nevertheless, the majority of the studies mentioned above (Table 1) were designed mainly on microsatellites, and focus on fewer loci and higher mutation rates than MPS data. In addition, studies centered on MPS markers were mostly based on bio-informatic simulations (Table 1). Among these, three studies have evaluated the effect of sampling design on estimates of genetic parameters using empirical and genomic data. Puckett & Eggert (2016) compared data for 15 microsatellite and 1000 SNPs in *Ursus americanus* and found that the SNP dataset was more precise than microsatellites to assign the provenance of 96 individuals belonging to 34 populations. Nazareno et al., (2017) analyzed different sampling tests for two populations of *Amphirrhox longifolia* (Violaceae), ~4,000 SNPs and 70 individuals. They found that sampling over 8 individuals and 1000 SNPs did not increase test accuracy. Flesch et al., (2018), analyzed four populations of rocky mountain bighorn sheep, 14,000 SNPs and 120 individuals in total, finding that the accurate estimation of genetic parameters was achieved after sampling 25 individuals per population.

While the studies of Puckett & Eggert (2016), Nazareno et al., (2017) and Flesch et al., (2018) are informative and relevant, they were performed in 34, 2, and 4 populations and they were based in a relatively small number of individuals (96, 70, 120) or SNPs (1000, ~4,000 and ~14,000, respectively). More importantly, these studies did not test the effect of sampling design on the detection of outlier SNPs using empirical data. As mentioned above, sampling few individuals or loci might affect the precision in the estimation of summary statistis, and sampling few populations could affect estimates of patterns of isolation, and the detection of outlier SNPs (de Mita et al. 2013). Therefore we aim at testing the effect of sampling design – increasing and varying the number of populations, number of individuals and number of SNPs-to assess with greater precision the potential biases and errors in estimates of population genomics parameters, in patterns of isolation and in the detection of outlier SNPs.

In this study, we analyzed two large genomic dataset (33,454 SNPs, 646 individuals and 49 populations obtained with the MaizeSNP50 Genotyping BeadChip; and 9,735 SNPs, and indivudals pooled from 47 populations obtained with the DArTseqTM data) and one microsatellite (22 microsatellite loci, 527 individuals and 29 populations) datasets of Mexican wild maize populations (*Zea mays* ssp. *mexicana* and *Zea mays* ssp. *parviglumis*) to explore the effects of sampling design in the estimation of population genomics parameters (*H*s, *F*_IS_, *F*_ST_), landscape genetics (tests of isolation by distance and environment), test of centrality (association between genetic diversity, ecological niche, and distance from the center of the distribution (Eckert et al., 2008; Lira-Noriega et al., 2013; Aguirre-Liguori et al., 2017)), and estimation of candidate SNPs (outlier SNP detection tests). In particular, we compared the effect of 1) using MPS vs. microsatellites makers; 2) using individual and data with known ascertainment bias (MaizeSNP50 Genotyping BeadChip) vs. pooled and non-ascertained biased data (DArTseqTM data); 3) varying the number of sampled loci (genomic datasets: 100, 1,000, 5,000, 15,000; microsatellite datasets: 5, 10, 15); 4) varying the number of sampled individuals per population (3, 6 and 9 individuals); 5) changing the number of sampled populations (5, 10, 20, 30, 40 populations); 6) varying the number of individuals per population; and 7) testing the effect of the number of sampled populations in the detection of outlier SNPs. Based on these results we propose guidelines for genomic sampling design in accordance to study objectives.

## MATERIALS AND METHODS

### Studied taxon

Mexican wild maize, or teosintes, are divided into two main subspecies, the lowland subspecies *Zea mays* ssp. *parviglumis* (hereafter *parviglumis*) and the highland subspecies *Zea mays* ssp. *mexicana* (hereafter *mexicana*) (Aguirre-Liguori et al., 2016). Cultivated maize was domesticated from *parviglumis* around 9,000 years ago (Matsuoka et al., 2002). Given the proximity of teosintes to maize, different genomic resources are available (Hufford et al., 2012; Aguirre-Liguori et al., 2016) and several studies have analyzed their population genomics (van Heerwaarden et al., 2010, 2011; Hufford et al., 2013; Pyhäjärvi et al., 2013; Aguirre-Liguori et al., 2017, 2019a,b; Fustier et al., 2017, 2019; Moreno-Letelier et al., 2018). Briefly, genomic studies suggest that teosintes have high genetic diversity, show patterns of isolation by distance and environment, and show strong patterns of local adaptation (Pyhäjärvi et al., 2013; Aguirre-Liguori et al., 2017, 2019a, b; Fustier et al., 2017, 2019). The vast genomic resources and biological knowledge makes teosintes an ideal system to study the importance of sampling design in analyses of genetic diversity, isolation patterns and identification of candidate SNPs, since sampling biases can be compared with the available knowledge.

### Datasets

First, we combined the MaizeSNP50 Genotyping BeadChip data published by Pyhäjärvi et al., (2013) and Aguirre-Liguori et al., (2017) to obtain a total dataset consisting of 49 populations, 24 belonging to *mexicana* and 25 to *parviglumis* (Supporting Information Figure S1), including between 12 and 15 individuals per population, and 33,454 SNPs. Since the

MaizeSNP50 Genotyping BeadChip was designed to maximize variation in maize, it has ascertainment bias (Alberchesten et al., 2010). Therefore this dataset is expected to sequence SNPs that are in high frequency across distant teosintes populations and might overestimate genetic diversity and underestimate genetic differentiation. We also downloaded the DArTseqTM data from Aguirre-Liguori et al., (2019a), which is composed of pooled DNA formed by ~12 individuals belonging to 47 populations, 21 belonging to *parviglumis* and 26 to *mexicana* (Supporting Information Figure S1) and 9,735 SNPs. The DTS dataset is based on initially cutting the DNA using restriction enzymes (Sansaloni et al., 2011; Ren et al., 2015) and has lower ascertainment bias (see Aguirre-Liguori et al., 2019a). This dataset has a non-biased frequency spectrum and is expected to generate more robust demographic inferences (Alberchesten et al., 2010). We called the Illumina and the Dartseq datasets the 50K and DTS datasets, respectively.

To be able to compare deviations obtained from MPS and microsatellite markers, which have been used in many studies (Table 1), we also downloaded the microsatellite dataset from Gasca-Pineda et al. (2019), which includes 527 individuals distributed across 29 populations, 14 belonging to *parviglumis* and 15 to *mexicana* (Supporting Information Figure S1), 22 loci and 355 alleles. We also downloaded the environmental and geographic information of each population, including the longitude and latitude at which they grow, as well as the score of the first principal component (PC1) describing temperature (Supporting information in Aguirre-Liguori et al., 2017, 2019a, Gasca-Pineda et al., 2019). The environmental data was obtained from 19 bioclimatic variables downloaded from WorldClim at a 30°arc Resolution, and the PCA was performed using the prcomp function in R across al variables and populations.

Importantly, these three datasets share many populations (Supporting Information Figure S1) and are distributed along the entire geographic and environmental distribution of teosintes (Hufford et al., 2012; Pyhäjärvi et al., 2013; Aguirre-Liguori et al., 2017, 2019a). They are composed of many individuals per population (between 9 and 26 individuals per population) and contain a large number of SNPs or microsatellite markers, distributed along the 10 chromosomes of teosinte. Therefore, we defined these datasets as the samples representing the most accurate data (i.e., the “real” data for the purpose of this work). We performed different subsamplings of number of loci, number of individuals per population and number of populations sampled to test deviations generated by different sampling designs on the estimation of population genetics parameters, landscape genetics and tests for local adaptation (see below for descriptions of the sub-samplings). For all subsamplings we combined the *mexicana* and *parviglumis* populations. We also tested the effect of sampling design separating by subspecies to account for taxon-bias. However since the patterns were similar between the entire datasets and the subspecies datasets, for simplicity we present the results of the combined dataset and show results of each subspecies in supporting information. We used *adegenet* (Jombart, 2008) and *hierfstat* (Goudet 2005) packages of R 3.0.2 (R Core team) to generate *genind, genpop* and *hierfstat* objects to manipulate the data. All these objects are indexable, and therefore allow subsampling random individuals, SNPs, microsatellite markers, subspecies, and/or populations.

### Estimation of population genetics parameters

For each entire dataset and each subsampling within dataset (see below for descriptions of the sub-samplings), we used the *basic.stats* function of the *hierfstat* package in R to calculate the sample *H*_S_ and *F*_IS_ and *F_ST_*. For the genomic datasets, we also used the *BEDASSLE* package (Bradburd et al., 2013) in R to calculate the pairwise *F*_ST_ between populations, according to Weir & Hill (2002). For the microsatellite datasets, we used the *pairwise.fst* function of the *hierfstat* package in R to calculate Nei’s pairwise *F*_ST_ between populations. For each summary statistic, we obtained the mean value across loci for each population.

We tested patterns of isolation by distance (IBD) and isolation by environment (IBE) using multiple regressions of distances matrices (*MRM* function from the *ecodist* package) to test the association between pairwise genetic distance (*F*_ST_) as a response variable and the environmental and geographic distances as predictive variables. We performed 1,000 permutations to test for significance between variables. The advantage of MRM tests is that it allows testing simultaneously both the environmental and geographic distances, and determines the relative contribution of each variable (Lichstein, 2007).

Finally, we tested the central abundance hypothesis (CAH), which suggests that genetic diversity should reduce as a function of the distance from the geographic or niche centroid (Eckert et al., 2010; Martínez-Meyer et al., 2012; Lira-Noriega & Manthey, 2013). For the CAH tests, we used simple linear regressions (*lm* function in R) to test the association between *H*s as response variable and the distance to the niche and geographic centroids as independent variables (which were estimated as the Euclidian distances from the geographic and niche centroids; for more details of the methods see Aguirre-Liguori et al., 2017).

### Sampling designs

#### subsampling of the number of loci and the number of individuals per population

First, we tested the effect of sampling different number of SNPs or microsatellite markers per population. We used R custom scripts (available as supporting information-function num_locs) to extract data from the entire sets: for the DTS dataset 100, 1000 and 5,000 random SNPs; for the 50K dataset 100, 1000 or 15,000 random SNPs; and for the microsatellite dataset 5, 10 and 15 random markers. For the 50K and microsatellite datasets, we also tested the effect of sampling different estimates of individuals per populations. We extracted randomly for each population 3, 6 and 9 individuals (available as supporting information-function num_inds()). This was not performed on the DTS dataset, because it is based on pooled DNA.

For the number of SNPs, number of microsatellite markers and number of individuals per population, we re-sampled randomly and without replacement each set 1000 times, we generated *genind/hierfstat/BEDASSLE* input objects and estimated the summary statistics described above (*H*s, F_IS_, *F*_ST_, IBE, IBD, CAH associations). For each parameter and each replicate we obtained the mean summary statistic across populations and generated a distribution based on 1000 summaries corresponding to each subsampling. We compared qualitatively these simulations to the “real” dataset, to determine the deviations generated by different sampling designs.

#### Subsampling the number of populations

To test the effect of the number of populations in the estimation of the parameters described above (*H*s, *F*_IS_, *F*_ST_, IBE, IBD, CAH associations), we performed random sampling designs. For the genomic datasets, we sampled 5, 10, 20, 30 and 40 random populations from the 49 (50K) and 47 (DTS) populations described above (Supporting Information Figure S1). For the microsatellite dataset, we sampled 5, 10 and 20 random populations from the 29 populations described above (Supporting Information Figure S1). Again, we generated 1,000 sub-samples without replacement (available as supporting information-function num_pops ()), and for each replicate we generated *genind/genpop/hierfstat/BEDASSLE* inputs and in each case, we calculated the estimates described above, and for summary statistics we estimated the mean across populations.

#### Tradeoff between number of individuals and populations

To test the tradeoff between the number of individuals and number of populations, we also tested three sets of sampling designs changing the number of inividuals sampled per population, going from few individuals in many populations to many individuals y few populations. For three scenarios (3 individuals and 49 populations; 6 individuals and 24 populations and 9 individuals and 10 populations) we generated 1,000 subsamples and estimated the paramteres described above.

### Test for local adaptation

Detecting outlier SNPs is challenging, since high genetic structure can inflate false positives (Schoville et al., 2012; de Mita et al., 2013; Tiffin & Ross-Ibarra, 2014). We tested the effect of varying the number of populations in detecting outlier loci. For this, we sub-sampled without replacement 5, 10, 20 and 30 random populations from the entire 50K dataset (49 populations and between 12-15 individuals per population, see Aguirre-Liguori et al., 2017 for more details). Since outlier analyses are time-consuming, we only generated 10 replicates of each sampling design, and we sub-sampled 10,000 SNPs from the 50K dataset. We also ran 10 times the analysis with the 49 populations to have comparable number of replicates. We chose 10,000 SNPs to reduce computing times and because our results (see below) show that over 1,000 SNPs are enough to obtain a reliable *F*_ST_ estimate among populations.

For each sample, we used *Bayescenv* (De Villemereuil & Gaggiotti, 2015) to identify outlier SNPs associated to PC1 (as in Aguirre-Liguori et al., 2017, 2019a). *Bayescenv* decomposes *F*_ST_ based on a signal shared between all loci (β), a signal specific to each locus

(α) and the association of the SNP with the environmental variable tested (γ). We used default parameters to run the analyses and we defined outlier SNPs as those that had *q-val* <0.05, which is a conservative approximation to detect outlier loci (de Villemereuil & Gaggiotti, 2015). For each replicate of each sampling design, we recorded the highest *F*_ST_ value for a SNP and the number of SNPs that had *q-val* <0.05.

## RESULTS

### Summary statistics for the entire datasets

We considered the entire datasets (50K, DTS and microsatellite) as those revealing the “real” or most accurate patterns of genetic diversity across teosinte populations. Table 2 shows the mean *H*_S_, *F*_IS_ and *F*_ST_ across populations, patterns of isolation by distance and environment, and test of central abundance, estimated for the DTS, 50K and microsatellite datasets (the distribution across different sampling designs are found Supporting Information Table S1).

**TABLE 2.** Summary statistics estimated for the DTS, 50K and microsatellite datasets of Mexican wild maize. For the mean and maximum and minimum values across 1000 replicates of each sampling designs see Supporting Information Table S1.

| Mean estimate | DTS | 50K | microsatellite |
| --- | --- | --- | --- |
| $H_S$ | 0.130 | 0.225 | 0.691 |
| $F_{IS}$ | | 0.069 | 0.182 |
| $F_{ST}$ | 0.393 | 0.246 | 0.106 |
| MRM: geographic ($\beta$) | 0.027 | 0.025 | 0.013 |
| MRM: environmental ($\beta$) | 0.011 | 0.011 | 0.004 |
| CAH: geographic ($\beta$) | -0.014 | -0.014 | -0.041 |
| CAH: environmental ($\beta$) | -0.006 | -0.008 | -0.031 |

We found striking differences between the datasets for the estimated mean across populations of *H*s, *F*_ST_ and *F*_IS_ (Figure 1, Table 2). We detected that DTS dataset presents low mean genetic diversity across populations (*H*s=0.13), the 50K intermediate values (*H*s=0.225) and the microsatellite data high values (*H*s=0.69). In parallel fashion, we found that DTS presents the highest mean genetic structure across pairwise comparisons (*F*_ST_=0.393), followed by the 50K dataset (*F*_ST_ =0.246) and finally the microsatellite dataset (*F*_ST_ =0.11). We were not able to calculate *F*_IS_ for the DTS dataset (pooled DNA), but we also found differences between the estimated mean using the 50K (*F*_IS_=0.065) and microsatellite datasets (*F*_IS_=0.19).

**FIGURE 1.**
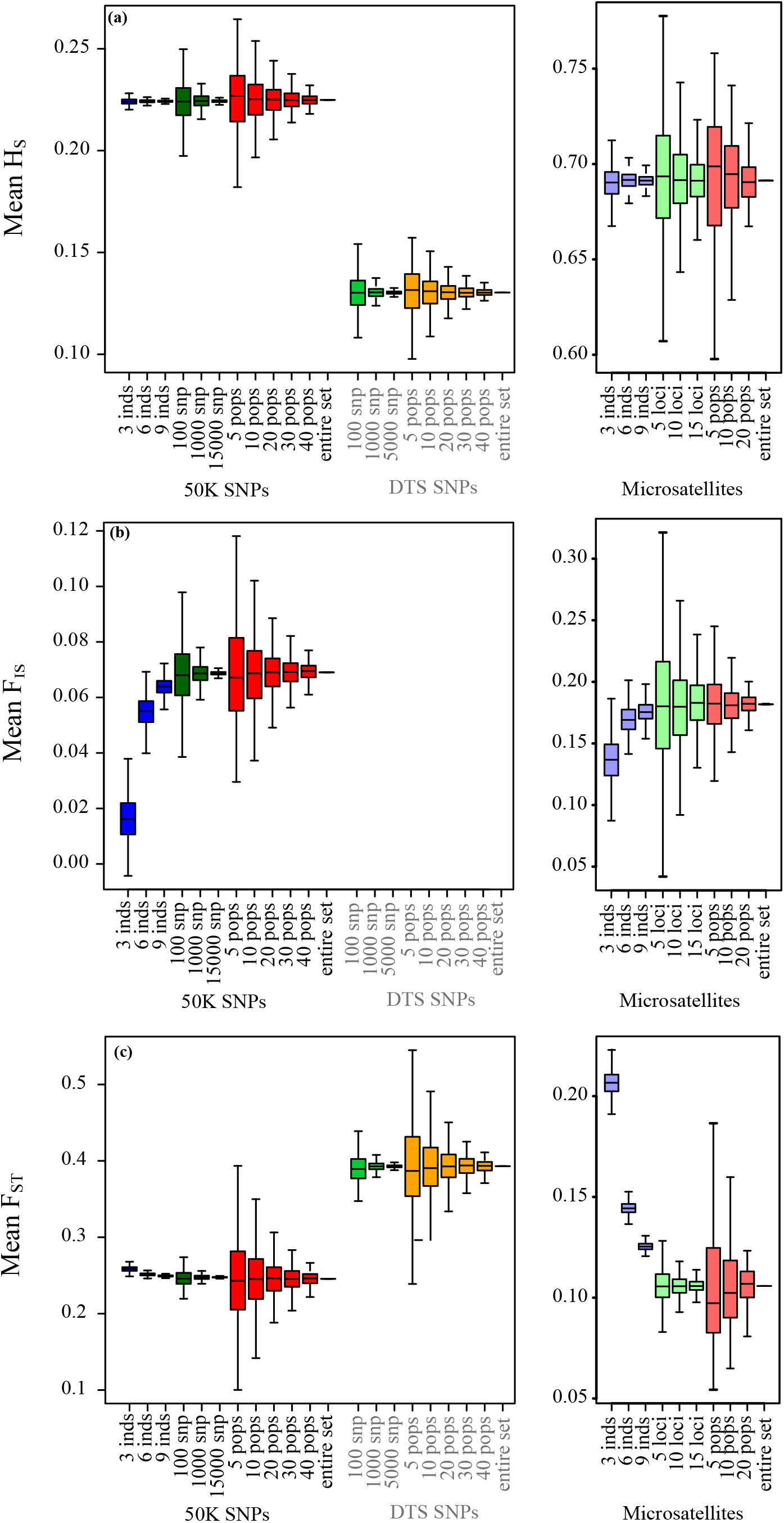
Effect of sampling designs on the estimation of summary statistics for genomic (left panels) and microsatellite (right panels) datasets: (a) *H*_S_; (b) *F*_IS_; (c) *F*_ST_. Boxplots show the distribution of mean summaries estimated for 1,000 replicate simulations varying the number of individuals, number of SNPs, and number of populations sampled. Significantly different distributions are shown in Supporting Information Table S2. *F*_IS_, was not possible to obtain for the DTS dataset because it is based on pooled data.

In contrast to the summary statistics, for the three datasets we found similar patterns of IBD and IBE (Figure 2), based on MRM tests (Figure 2, Table 2). For the three datasets, we observed that patterns of IBD and IBE were positive and that the values were stronger for IBD than IBE. Finally, for the three datasets, we observed negative associations between genetic diversity and the distance to the geographic and niche centroids (Figure 3, Table 2).

**FIGURE 2.**
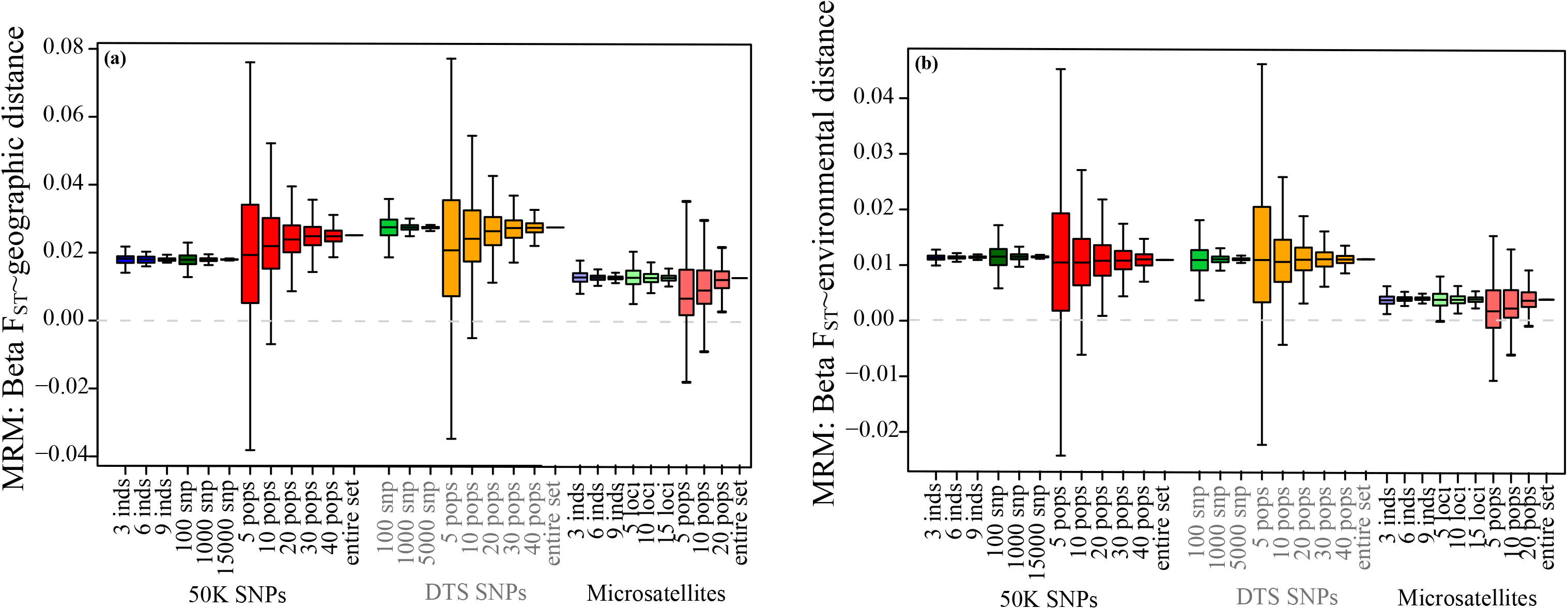
Effect of sampling designs on the analysis of patterns of isolation for genomic and microsatellite datasets: (a) IBD-MRM test; (b) IBE-MRM test. Boxplots show the distribution of associations estimated for 1,000 simulations varying the number of individuals, number of SNPs, and number of sampled populations. The dotted grey line shows the 0 value. A list of significantly different distributions is shown in Supporting Information Table S2.

### Varying the number of sampled individuals

As mentioned above, this test was only possible to perform with the 50K and microsatellite datasets, since they were based on individual samples.

When sampling different estimates of individuals, we found that the 50K and microsatellite datasets were not sensitive to estimates of *H*s (Figure 1a, Supporting Information Fig. S2), patterns of isolation by distance and environment (Figure 2, Supporting Information Fig. S3) and tests for association between genetic diversity and ecological variables (Figure 3, Supporting Information Fig. S4). In fact, the range of summary estimates generated by sampling fewer individuals, fell within the range of values obtained when sampling 30 populations of the 50K dataset and 20 populations of the microsatellite dataset (Compare boxplots in Figures 1-3; Supporting Information Table S1, S2). While the ranges of the maximum and minimum values were close to the “real” estimates for the 50K dataset, many of the comparisons between subsamples of 3, 6 and 9 individuals and the real data set were nonetheless significantly different (Supporting Information Table S2-S4).

**FIGURE 3.**
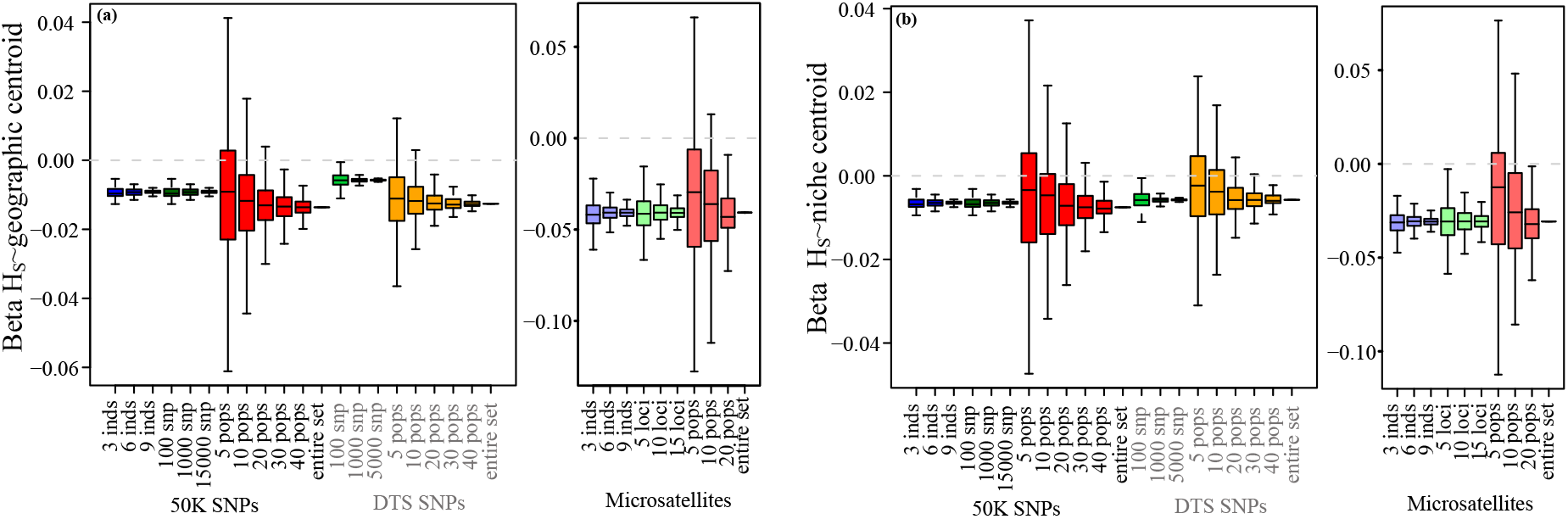
Effect of sampling designs on the estimation of the central abundance hypothesis for genomic and microsatellite datasets: (a) Association between distance to the geographic centroid and Hs; (b) Association between distance to the niche centroid and Hs. Boxplots show the distribution of associations estimated for 1,000 simulations varying the number of individuals, number of SNPs, and number of sampled populations. The dotted grey line shows the 0 value. A list of significantly different distributions is shown in Supporting Information Table S2.

In contrast, changing the number of sampled individuals generated strong deviations for the estimation of *F*_IS_ for the 50K and microsatellite datasets (Figure 1b, Supporting Information Fig. S2), moderate deviations for estimates of *F*_ST_ for the 50K dataset (Figure 1c, Supporting Information Fig. S2, Table S1, S2) and very strong deviations for the estimates of *F*_ST_ for the microsatellite dataset (Figure 1c, Supporting Information Fig. S2). Sampling fewer individuals reduced *F*_IS_ estimates and increased *F*_ST_ estimates (Figure 1b,c).

### Varying the number of sampled loci

The three datasets showed similar responses to sampling different number of loci. In all cases, sampling fewer loci increased the variance across replicates for all estimates (Figures 1-3; Supporting Information Fig. S2-S4). Importantly, we found that the variance across replicates was higher for the microsatellite dataset, followed by the DTS dataset and finally when sampling each subspecies using the 50K dataset (Supporting Information Fig. S2-S4).

Even though decreasing the number of loci increased the variance, it was interesting to note that estimated distributions fell close to the estimates for “real” datasets (Supporting Information Table S1), and that the distribution of different estimates fell within the range of 30 sampled populations using all loci and all individuals for the DTS and 50K datasets (compare Boxplots ranges in Figures 1-3). For the microsatellite dataset we found that sampling fewer loci increased the variance within the distribution of 20 sampled populations (compare ranges) using all loci and all individuals for *F*_ST_ (Figure 1c), the tests for IBD and IBE (Figure 2) and the test of the central abundance hypothesis (Figure 3). For *H*_S_ and *F*_IS_, sampling fewer microsatellite loci produced an important increase in the variance across replicates.

The only test that generated strong deviations with respect to the “real” datasets (even if the ranges were not very large) was the association between *H*_S_ and the geographic centroid, where we found that sampling fewer DTS and 50K SNPs reduced the association (β got closer to 0). Importantly, sampling 1,000 or 15,000 of the 50K SNPs; 1,000 or 5,000 of the DTS SNPs; and 10 or 15 loci of the microsatellite dataset generated significant similar summary statistics (Supporting Information Table S2).

### Varying the number of sampled populations

Varying the number of populations generated similar mean values to those found for the “real” datasets. However, for the three datasets, sampling 5 and 10 populations generated significant deviation from the real estimated values for isolation patterns and tests of centrality (Supporting Information Table S2-S4). The variance across datasets dropped after around 30 populations for the genomic datasets (see ranges in Supporting Information Table S1), but remained high for the microsatellite dataset.

Importantly, sampling different number of populations generated a high variance in the estimation of all parameters, except for *F*_IS_ and *F*_ST_ for the microsatellite dataset (Figure 1b,d) and *F*_IS_ estimates for the 50K dataset (Figure 1, Supporting Information Fig. S2-S4). In these cases, the deviations across replicates when sampling fewer populations was lower than the variance generated by sampling different number of individuals (Figure 1).

For patterns of IBD and IBE, we also found that sampling fewer than 10 populations generated in some replicates incorrect association estimates. The real value showed positive associations (Table 2), but for 5 and 10 sampled populations we found that up to 18% and 7% of sample replicates generated negative associations, respectively (Supporting Information Table S5; see changes in signs in Supporting Information Table S1). For tests of association between *H*_S_ and ecological variables, this was even more sensitive for the genomic datasets, since we found up to 4.6% of positive estimates (when the entire dataset was negative) for 30 populations when testing association between *H*s and the distance to the niche centroid. However, we found that the DTS was less sensitive to deviation for associations between ecological and genetic variables (Figure 3).

Finally, we found that the microsatellite dataset was more sensitive for isolation patterns (we found up to 30% of samples showed opposing patterns to the entire dataset when sampling 5 populations; Supporting Information Table S5), and less sensitive for tests of central abundance hypothesis (we found up to 1.8% of false positives when sampling 20 populations; Supporting Information Table S5).

### Tradeoff between number of individuals and number of populations

For the 50K dataset, we also contrasted the effect of sampling few individuals in many populations and many individuals in fewer populations (3 individuals/49 populations, 6 individuals in 24 populations, and 9 individuals in 10 populations). For all summary statistics, except FIS, we found that sampling more populations but fewer individuals generated more accurate results and lower biases (Figure 4; Supporting Information Fig. S5).

**FIGURE 4.**
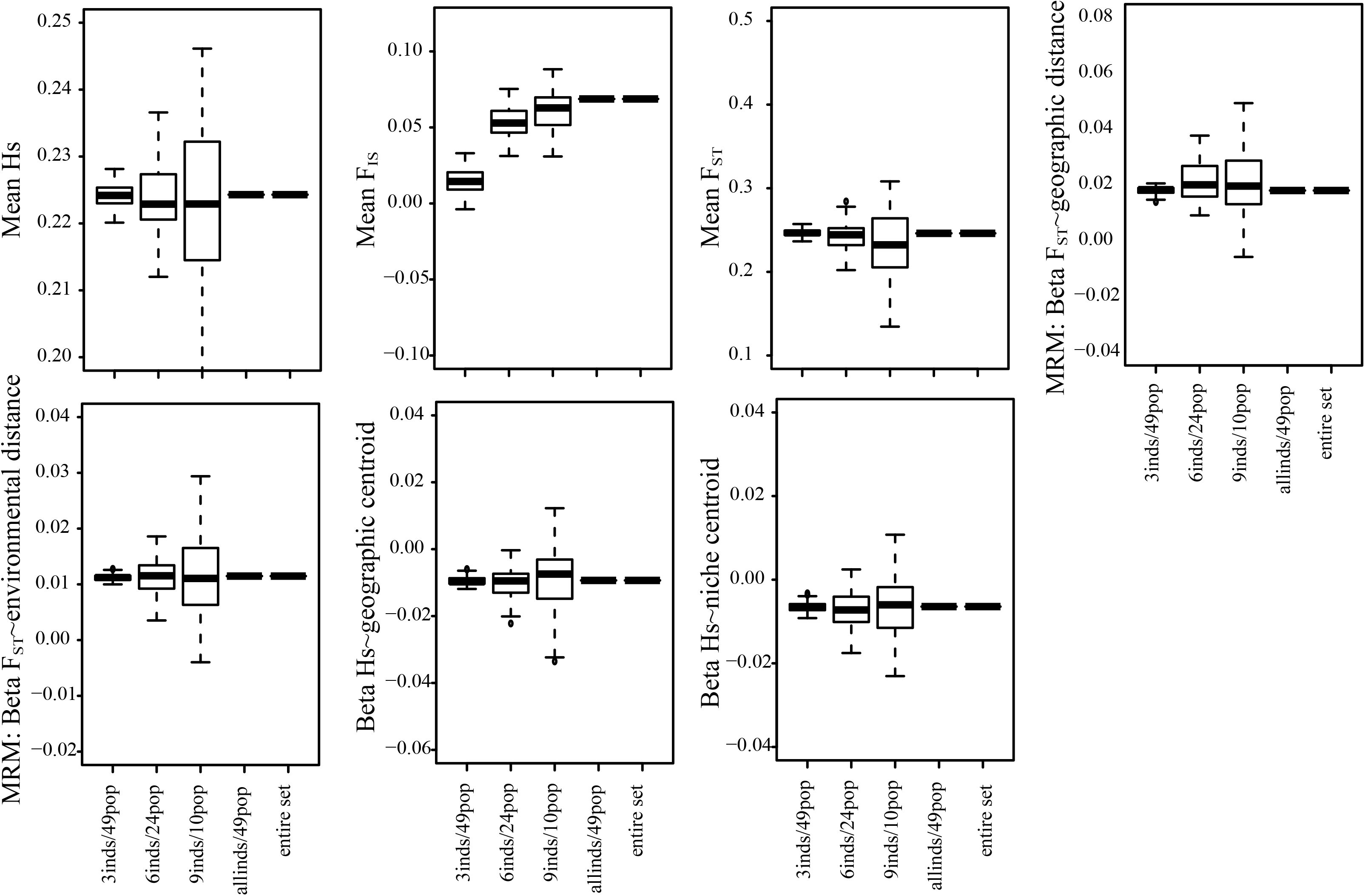
Trade off between the number of individuals and the number of populations sampled for all summary statistics using the 50K dataset. We tested the effect of sampling more individuals in few populations and few individuals in many populations.

### Varying the number of populations in the identification of candidate SNPs

Reducing the number of sampled populations increased the maximum *F*_ST_ value for a SNP. Figure 5a shows that for 5 and 10 sampled populations, the maximum *F*_ST_ for a locus found by *bayescenv* across replicates was significantly higher than for the rest of the sampling designs. We also tested the number of candidate SNPs across replicates. We found that more candidate SNPs when sampling a higher number of populations (Figure 5b). Interestingly, we found that few outlier SNPs were shared between replicates and between sample designs sampling designs (Supporting Information Table S6).

**FIGURE 5.**
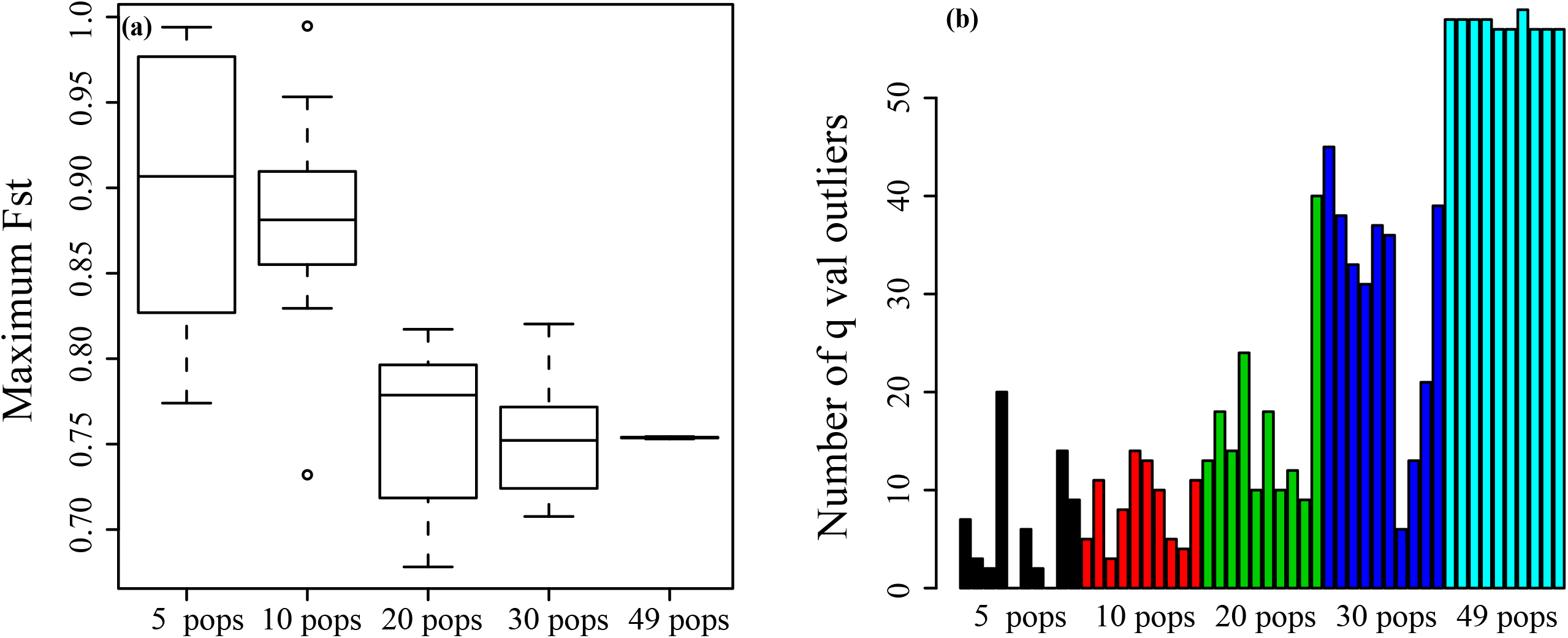
Effect of sampling different number of populations on the identification of outlier SNPs. (a) Distribution of the highest *F*_ST_ identified for a locus across simulations; (b) Number of outlier SNPs (*q-val)* identified for each replicate for different number of sampled populations.

## DISCUSSION

Massive parallel sequencing is allowing ecologists to understand better the evolutionary ecology of populations and understand the basis of adaptation (Stapley et al., 2010; Ross-Ibarra & Tiffin, 2014). Unfortunately, it remains challenging for many researchers to generate large genomic samples, posing a tradeoff between the information obtained with MPS and the number of populations sampled (Meirmans, 2015). Here we used two empirical genomic datasets and a microsatellite dataset obtained for a large sample of wild maize, the teosintes (*Zea maiz* ssp. *parviglumis* and *Zea mays* ssp. *mexicana),* to analyze the deviations generated by different sampling designs while estimating classic population genomics and landscape genomics estimates.

### Comparing datasets

We found that the most evident differences between datasets were associated to *H*_S_, *F*_ST_ and *F*_IS_ estimates. These differences are expected given the properties of each dataset. First, we found that the microsatellites had the highest *H*_S_ and lowest *F*_ST_, which is a well-known pattern, and can be explained by their large number of alleles and mutation rates (Ellegren, 2004). Second, the 50K dataset had higher genetic diversity and lower *F*_ST_ compared to the DTS dataset. This is concordant with the design of the 50K dataset to detect highly polymorphic SNPs in maize, and therefore has ascertainment bias (Alberchesten et al., 2010). For this dataset we expected higher genetic diversity and more shared polymorphisms. In contrast, the DTS dataset was generated using restriction enzymes (similar to GBS and other reduced representation methods, including RADtags), and has lower ascertainment bias (Sansaloni et al., 2011; Ren et al., 2015). Therefore, we expected to find lower genetic diversity and have a higher proportion of fixed alleles (Aguirre-Liguori et al., 2019a), increasing the *F*_ST_ estimate.

Even though we found differences between datasets, it was interesting to note that for tests of isolation, and tests associated to the central abundance hypothesis, very similar patterns were found with the three datasets. In all cases, we found the same associations for the summary statistics (Table 2). This is an interesting observation, and suggests that for these types of analyses any genomic or microsatellite dataset might be useful, as long as enough individuals, loci and populations are sampled (Table 3, See below).

**TABLE 3.** Recommendations for sampling designs depending on study objectives.

| Estimate | Number of individuals | Number of loci | Number of populations | Considerations |
| --- | --- | --- | --- | --- |
| $H_S$ | Not sensitive (>6 individuals) | Sensitive<br>(>1000 SNP loci; >15<br>microsatellite loci) | Sensitive (> 20<br>populations) | Increase the number of loci and populations. Genomic dataset is less<br>sensitive than microsatellite dataset |
| $F_{IS}$ | Very sensitive<br>(>9 individuals) | Sensitive<br>(>1000 SNP loci; >15<br>microsatellite loci) | Sensitive (>20 populations) | Increase the number of individuals over loci and populations. If a few<br>population are available, increase the number of individuals in those<br>populations |
| $F_{ST}$ | Microsatellite dataset was very<br>sensitive (>20 individuals)<br>50K dataset: Not sensitive (>9<br>individuals) | Not very sensitive<br>(>1,000 SNPs; >15<br>loci) | Sensitive (> 20<br>populations) | Increase the number of populations over the number of SNPs or<br>individuals |
| IBD and IBE<br>MRM tests | Not sensitive (>3 individuals) | Not sensitive<br>(>1,000 SNPs, >15<br>loci) | Very sensitive (>20<br>populations) | Sample as many populations as possible even if fewer individuals or<br>loci are sampled |
| IBD and IBE | Little sensitive (> 6<br>individuals) | Sensitive<br>(>1,000 SNPs; >15<br>loci) | Very sensitive (>20<br>populations) | Increase all samples. |
| CAH tests | Not very sensitive (>3<br>individuals for genomic<br>datasets; >6 individuals for<br>microsatellite datasets) | Sensitive depending<br>on the dataset<br>(>1,000 DTS SNPs,<br>>100 50K SNPs, >15<br>microsatellite loci) | Very sensitive (>30<br>populations) | Increasing the number of populations is more important than increasing<br>the number of loci or individuals. Microsatellites are more sensitive<br>than genomic datasets to the number of loci and individuals, although<br>less sensitive to the number of populations sampled |
| Tests of selection | Not tested | Sensitive (as many as<br>possible) | Very sensitive (>30-40<br>populations) | As many SNPs as possible are needed to differentiate outlier loci, also<br>to increase the probability of finding loci within selective regions.<br>Increase as much as possible the number of populations, covering the<br>largest geographic and environmental distribution.<br>A possibility is to use pooled-sample DNA. |

Finally, when we compared the sampling designs with their respective “real” datasets, we found that the DTS dataset generated fewer significant deviations for sampling designs than the 50K datasets (Supporting Information Table S4). While this is in part attributable to the fact that for the DTS dataset we were not able to test *F*_IS_ values, or probably the deviations generated by sampling different number of individuals, for some comparisons the DTS dataset showed less deviations. For instance, we found that the DTS dataset generated fewer significant deviations for sampling designs (Supporting Information Table S3) and that this dataset was more robust for patterns associated to ecological variables and patterns of isolation (Supporting Information Table S4). The robustness of the DTS could be attributable to the lower ascertainment bias and a non-biased site frequency spectrum (Aguirre-Liguori et al., 2019a).

### Sampling different number of loci

For the three datasets, we tested the effect of sampling different number of SNPs and different number of microsatellite markers. While many comparisons were significantly different with respect to the “real” datasets (Supporting Information Table S4), sampling different number of SNPs did not produced large deviations from real statistics (Figure 1-3, Supporting Information Fig. S2-S4) for the genomic datasets, as long as 1000 SNPs or more were sampled. Sampling fewer microsatellite markers, increased slightly the variance across replicates for summary statistics (Figure 1) and tests of the CAH (Figure 3).

In general, it has been suggested that if a few populations and individuals are sampled, increasing the number of SNPs can increase the accuracy of estimates (See summary and references in Table 1). We did not test for a combination of sampling at the same time different number of SNPs and different number of populations, but we observed that when many populations are sampled (>20), after increasing the number of SNPs from 1,000, it produces similar patterns as sampling 5,000 (DTS) or 15,000 (50K) SNPs (Supporting Information Table S2). These are interesting observations, since depending on the study, it may be convenient to reduce genome or microsatellite coverage to increase the number of sampled populations, especially when patterns of isolation and demographic history are analyzed. However, it is important to notice that if the objective is to find targets of selection then, increasing the number of SNPs is critical to detect stronger neutral expectations and reduce false positives (de Mita et al., 2013), as well as increase the probability of finding SNPs that fall within coding or regulating regions (Glenn, 2011; Ekblom & Galindo, 2004).

### Sampling different number of individuals

We were only able to compare the effect of sampling different number of individuals for the 50K and microsatellite datasets. In general we found that for the 50K and microsatellite datasets many comparisons, especially for summary statistics, between the “real” dataset and the sampling designs were significantly different (Supporting Information Table S4).

However, for the 50K dataset we found that the majority of summary statistics distributions (except *F*_IS_) had a low variance that fell within the distribution of sampling 20-30 populations (Figures 1-3, Supporting Information Fig. S2-S4), indicating that sampling few individuals would not deviate summary statistic estimates. For the microsatellite dataset, we found that sampling 3-6 individuals generated important deviations for *F*_IS_ and *F*_ST_ estimates (Figure 1), and larger variance than the 50K dataset across replicates in all tests. Importantly, we found that sampling few individuals did not produce important deviations for tests of patterns of isolation, especially when using MRM tests (Figure 2).

For the 50K and microsatellite datasets, sampling fewer individuals increased the variation across sampling, but more importantly underestimated the *F*_IS_ inbreeding estimation (Figure 1b, Supporting Information Fig. S2). In contrast to other summary statistics, *F*_IS_ was very sensitive to the number of individuals sampled for both the microsatellite and 50K datasets. To identify what could be generating this differences, we correlated the mean FIS across populations and the number of missing data per sampling. We found that the reduction in F_IS_ correlated with increased NA associated to lower sampling (Supporting information Fig. S6).

These results suggest that for genomic datasets, as long as many populations are sampled, and *H*s, *F*_ST_, patterns of isolation, or patterns associated to ecological variables are tested, the number of individuals is not as sensitive as the number of populations sampled covering a large portion of the distribution (Table 3). In fact we found that it is more convenient to sample few individuals in as many populations as possible than the opposite (Figure 4, Supporting Information Fig. S5). On the contrary, studies that depend on *F*_IS_ values (i.e., genetic analyses in conservation studies), or that are performed using microsatellite data, should sample as many individuals as possible to reduce the bias generated by missing data (See summary and references in Table 1, Supporting information Fig. S6). Finally, for both datasets, our results corroborate studies that have shown that as long as many populations and enough loci are sampled, few individuals (Figure 4, Table 3) are needed to estimate patterns of isolation (See summary and references in Table 1).

### Sampling different number of populations

We tested the effect of randomly sampling different number of populations for all datasets. Sampling above 10 populations did not generate significant differences between sampling designs and the “real” sample for the three datasets (Supporting Information Table S4). However, we found that the number of populations was strongly associated with the accuracy of the mean estimates across replicates (Figures 1-3; Supporting Information Fig. S2-S4, Table S1). Sampling different number of populations with the microsatellite dataset generated a lower variance across replicates than the genomic datasets when estimating patterns of isolation using the MRM test (Figure 2a,b).

Importantly, we found that sampling few populations in some cases can result in opposite associations compared to the real dataset for patterns of isolation and ecological associations (i.e., less than 10 populations for patterns of isolation, and less than 30 populations for ecological associations). Although these incorrect associations were recorded only for a few replicates (Supporting Information Table S5), it is important to notice, that an overestimation of false associations could result by not sampling the entire geographic and environmental distribution (Rico, 2017; Chao et al., 2014).

The fact that few populations increases variance across replicates of genomic datasets is important, because many genomic studies usually sample few populations in order to increase the genomic coverage (Meirmans, 2015). Our results are concordant with different studies performing simulations that have shown that increasing the number of populations increases the accuracy in estimates of summary statistics and especially in landscape genetics studies (See summary and references in Table 1). In fact we found that it is more convenient to sample more populations with few individuals than few populations with many individuals (Figure 4, Supporting Information Fig. S5).

### Sampling different number of populations for detecting outlier SNPs

An important advantage of MPS is that it allows detecting candidate loci under selection. However, an important limitation of genome wide association studies is that demographic history and complex genetic structure can increase false positives (de Mita et al., 2013; Ross-Ibarra &Tiffin 2014; Schoville et al., 2013). Since adaptive loci could be important for conservation (Allendorf et al., 2010) and to respond to environmental change (Exposito-Alonso et al., 2017; Bay et al., 2018; Aguirre-Liguori et al., 2019b), many efforts have been made to reduce false positives and to better-detect genes that could be under selection.

We evaluated the effect of sampling different number of populations for detecting candidate SNPs. When we sampled few populations, it was interesting to notice that mean and maximum *F*_ST_ values across loci were significantly higher, supporting that sampling few populations could increase false positives or produce inaccurate estimates of real *F*_ST_ patterns (Figure 5a). A higher *F*_ST_ between populations complicates the accurate estimation of outliers (de Mita et al., 2013). Interestingly, we found that as more populations were sampled and F_ST_ was estimated more accurately, more outlier SNPs were identified (Figure 5b).

From these analyses, we conclude that increasing the number of populations is very important for detecting candidate SNPs since it allows to define more accurately the genetic structure (de Mita et al., 2013); and that it is important to cover the largest geographic and environmental distribution. These are important observations especially when not many populations can be sampled either because organisms have limited distributions (Smith & Wang, 2014; Chao et al., 2014) or because there is a tradeoff between the amount of SNPs that can be obtained using MPS and the number of populations that can be genotyped (Meirmans, 2015).

### Genomic sampling designs depending on the objectives

As mentioned before, there is a tradeoff between the amount of data that can be generated using MPS, and the number of populations sampled (Meirmans, 2015). The size of a dataset depends on many factors. First, there is a limit on the number of populations and individuals that can be genotyped. Some populations might be at risk, or few populations might exist or might be difficult to sample (Smith & Wang, 2014; Chao et al., 2014). Second, the number of SNPs called can depend on the sequencing platform (Metzker, 2010; Ekblom & Galindo, 2011; Glenn, 2011), the lab resources, the complexity of the genome (e.g., maize), and the availability of a reference genome (Metzker, 2010; Ekblom & Galindo, 2011; Glenn, 2011). Depending on the objectives, and the amount of data that can be produced using genomic platforms, we propose some suggestions for sampling design that could be considered according to our results (Table 3). It is important to consider that these recommendations are more reliable for species with life history traits similar to Mexican wild maize, and caution should be taken since life history can have an important effect in summary statistics (Hamrick & Godt, 1996; Nybom, 2004) and therefore on outlier tests.

*F*_IS_ is a measure associated to inbreeding (Wright, 1949; Allendorf et al., 2010; Waples, 2014), thus for studies of conservation genetics we propose that increasing the number of sampled individuals within populations is important for genomic and microsatellite datasets. If testing local adaptation is not a priority, then sampling fewer populations, but with many individuals (>20) might be more important, and with special focus on populations that could be at risk.

When testing summary statistics associated to demographic history, genetic structure, environmental associations, and landscape genomics, we found that the number of SNPs, and number of individuals did not produce strong deviations in comparison to the real data when testing the genomic datasets (even if some summary statistics were significantly different to the real data). However, we found that sampling few populations could create important deviations relative to the “real” summary statistics. In some replicates, we even found incorrect estimates of parameters (Figure 2, 3, 4; Supporting Information Fig. S2-S4; Table S5). These results corroborate the findings from other studies performed with microsatellites and using bio-informatic simulations (See summary and references in Table 1). Considering these results, we propose that if detecting local adaptation is not an objective and *F*_IS_ is not being measured, it is more important to sample many populations (~30) even if few individuals per population are considered (Figure 4) and few SNPs are obtained. Interestingly, we also found that DTS was less sensitive than the 50K dataset for different sampling designs (fewer significant deviations from real data, Supporting Information Table S4), and this might be explained by the reduced ascertainment bias that could increase accuracy (Alberchesten et al., 2010).

For studies whose aim is to identify candidate SNPs, we found that the number of sampled populations is critical, since sampling few populations could result in a lower detection power and an increase in false positives. Besides the number of sampled populations, it is important to also consider that the number of SNPs is crucial, since a higher number of SNPs will be more likely to fall within relevant genes or regions (Glenn, 2011; Ekblom & Galindo, 2011). If the number of samples that can be sequenced is an issue, we consider that it might be more convenient to increase the number of populations (>30) and SNPs by reducing the number of individuals per populations. Methods such as *Bayescenv* (De Villemereuil & Gaggioti, 2015), *Bayescan* (Foll & Gaggiotti, 2008) and *Bayenv* (Coop et al., 2010) do not rely on genotype counts, but rather on allelic counts. Therefore they are not sensitive to correct estimates of *F*_IS_ and one alternative can be to use a pooled sample approach. Aguirre-Liguori et al., 2019b used >33,000 50K individual based SNPs and ~9,000 DTS pooled based SNPs to test patterns of local adaptation across 7 environmental variables and although different number of SNPs were identified, the patterns were similar, suggesting that pooled data do not show significant bias in patterns of local adaptation and is a reliable alternative when this is the main objective of our study.

Finally, the microsatellite and the genomic dataset produced similar deviations when subsampling different number of individuals, loci and populations. In fact, for tests of patterns of isolation, the microsatellites showed fewer deviations for population sampling. Therefore, as long as enough individuals and loci are sampled, accurate estimates of genetic diversity are obtained with microsatellites. However, it might be more cost effective to use genomic datsets for those species for which microsatellites have not been developed.

## ACKNOWLEDGMENTS

We thank Yocelyn Gutiérrez Guerrero and Alberto Villasante Barahona for support in generating the microsatellite datasets. We thank Sarah Hearne and CIMMYT for generating the DTS dataset. We thank Erika Aguirre Planter and Laura Espinosa Asuar for technical support. Finally, we thank Gabriela Castellanos-Morales and Erika Aguirre-Planter for comments to the manuscript. JAAL thanks the “Programa de Doctorado en Ciencias Biomédicas, UNAM” and the scholarship provided by CONACYT (grant no. 255770). This work was funded by grants CB2011/167826 (CONACYT Investigación Científica Básica), CN-10-393 (UC MEXUS-CONACYT), and M12-A03 ECOS Nord France -CONACYT-ANUIES.

## DATA

The 50K dataset analyzed in this manuscript were downloaded from Dryad Digital Repositories: https://doi.org/10.5061/dryad.8m648 and https://doi.org/10.5061/dryad.tf556. the DTS dataset was download from CIMMYT Seeds of Discovery Dataverse under the accessibility link: http://hdl.handle.net/11529/10548175. The microsatellite dataset was downloaded from Supplementary Table S3 of Gasca-Pineda et al. 2019 (https://www.biorxiv.org/content/10.1101/820126v1.supplementary-material). R code used to perform different sampling designs are submitted as supporting information to this manuscript.

## AUTORS CONTRIBUTIONS

JAAL and LEE conceived and designed the work; JAAL and JALS performed and analyzed the genomic analyses; JGP generated, performed and analyzed the microsatellite datasets; All authors analyzed and interpreted the combined data. JAAL and JALS wrote the first version of the manuscript; All authors reviewed the manuscript.

