## Supporting information for "Evaluation of the minimum sampling design for population genomic and microsatellite studies. An analysis based on wild maize"

**Supporting Figures**

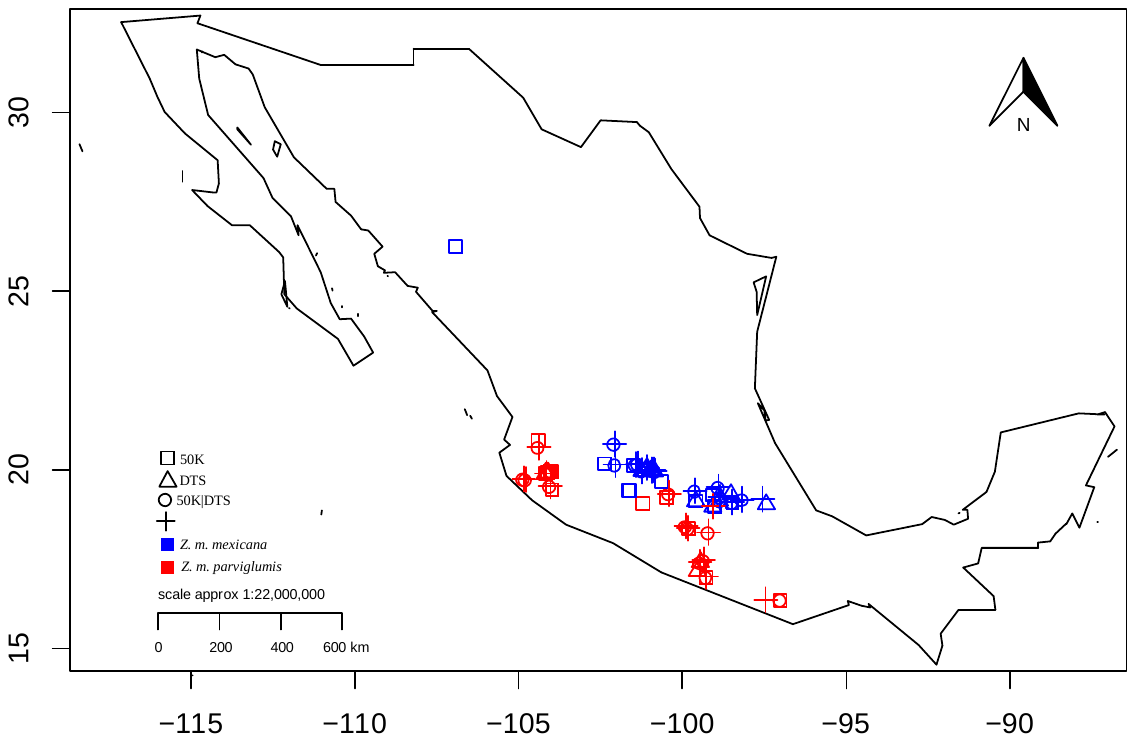

**Fig. S1** Map of the distribution of the studied populations.

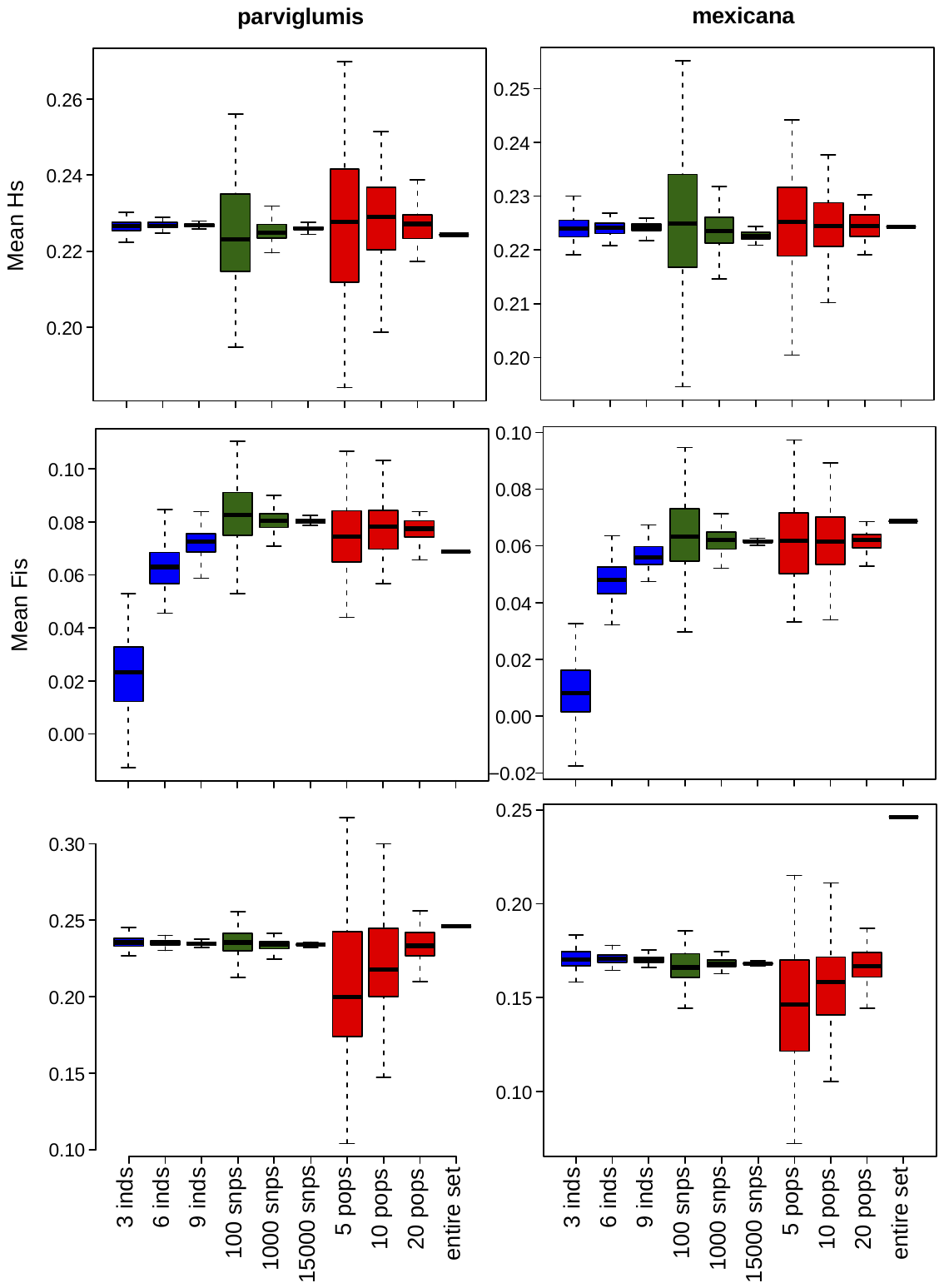

**FIG S2** Effect of sampling designs on the estimation of summary statistics for *Zea mays* ssp *parviglumis* and *Zea mays* ssp *mexicana* 50K genomic datasets: (above) *H*_S_; (intermediate) *F*_IS_; (below) *F*_ST_. Boxplots show the distribution of mean summaries estimated for 1,000 replicate simulations varying the number of individuals, number of SNPs, and number of populations sampled.

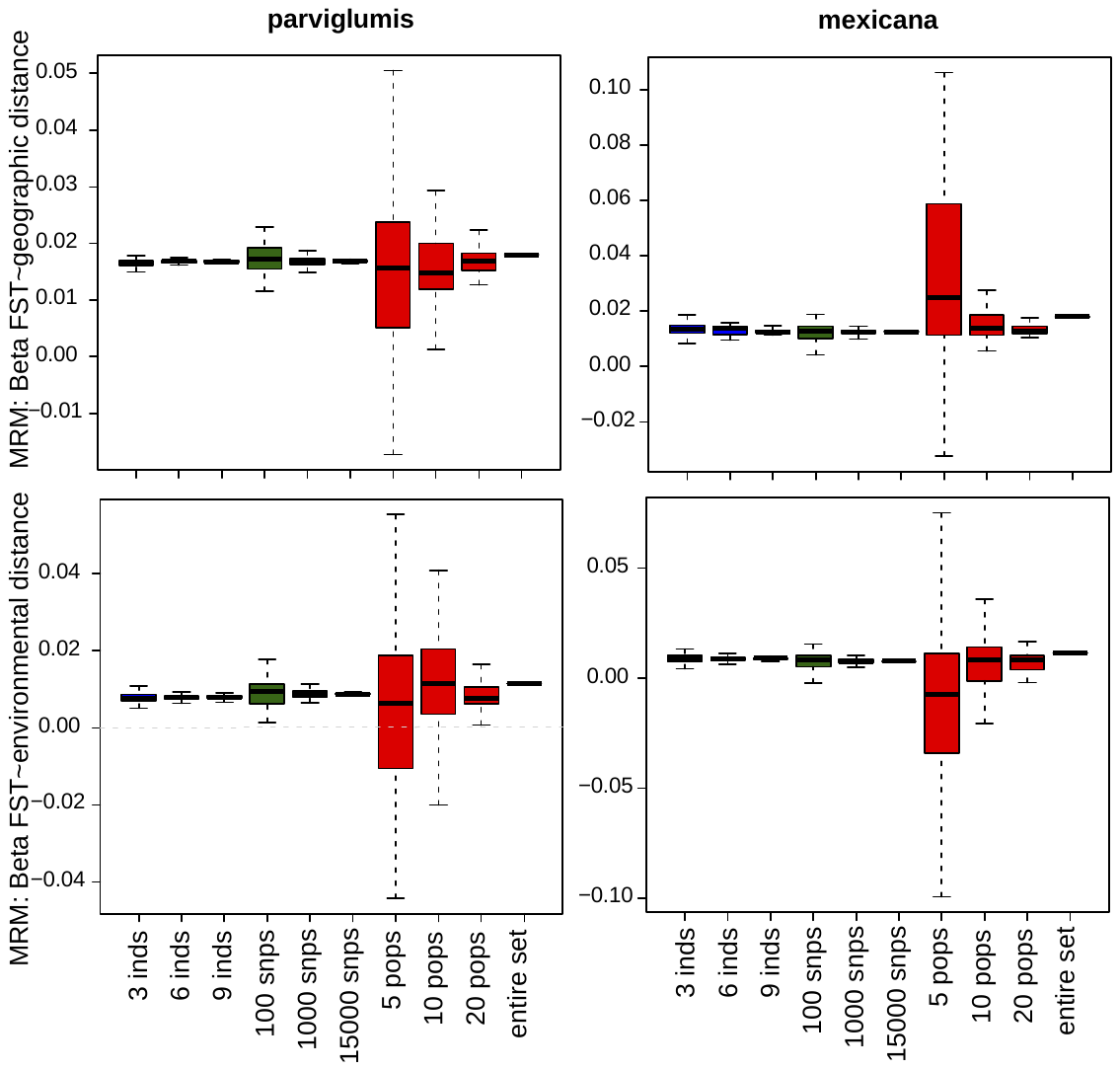

**FIG S3** Effect of sampling designs on the analysis of patterns of isolation for *Zea mays* ssp *parviglumis* and *Zea mays* ssp *mexicana* 50K genomic dataset: (above) IBD-MRM test; (below) IBE-MRM test. Boxplots show the distribution of associations estimated for 1,000 simulations varying the number of individuals, number of SNPs, and number of sampled populations. The dotted grey line shows the 0 value.

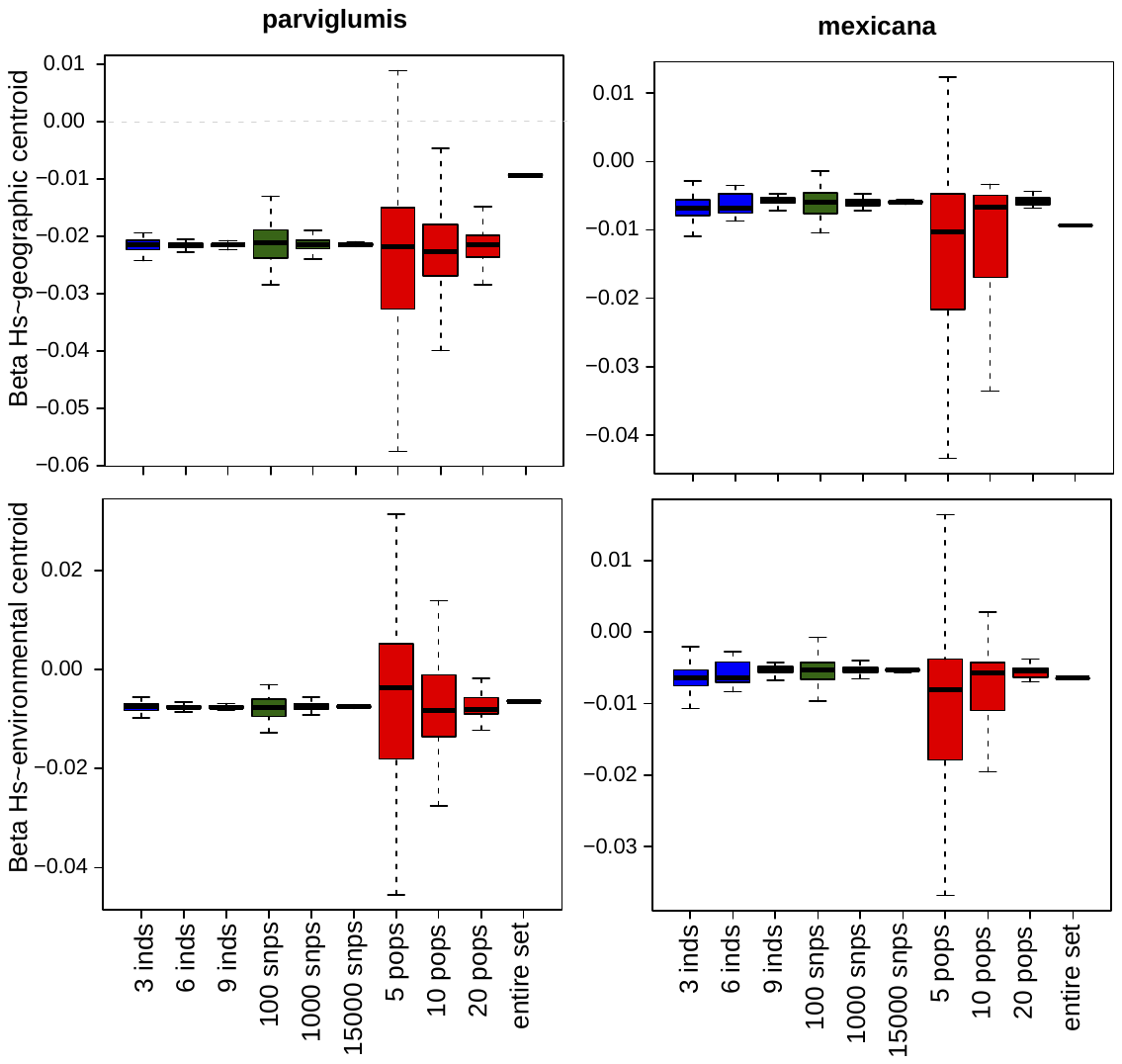

**FIG S4** Effect of sampling designs on the estimation of the central abundance hypothesis for *Zea mays* ssp *parviglumis* and *Zea mays* ssp *mexicana* 50K genomic dataset: (above) Association between distance to the geographic centroid and *H*s; (below) Association between distance to the niche centroid and *H*s. Boxplots show the distribution of associations estimated for 1,000 simulations varying the number of individuals, number of SNPs, and number of sampled populations. The dotted grey line shows the 0 value. A list of significantly different distributions is shown in Supporting Information Table S2.

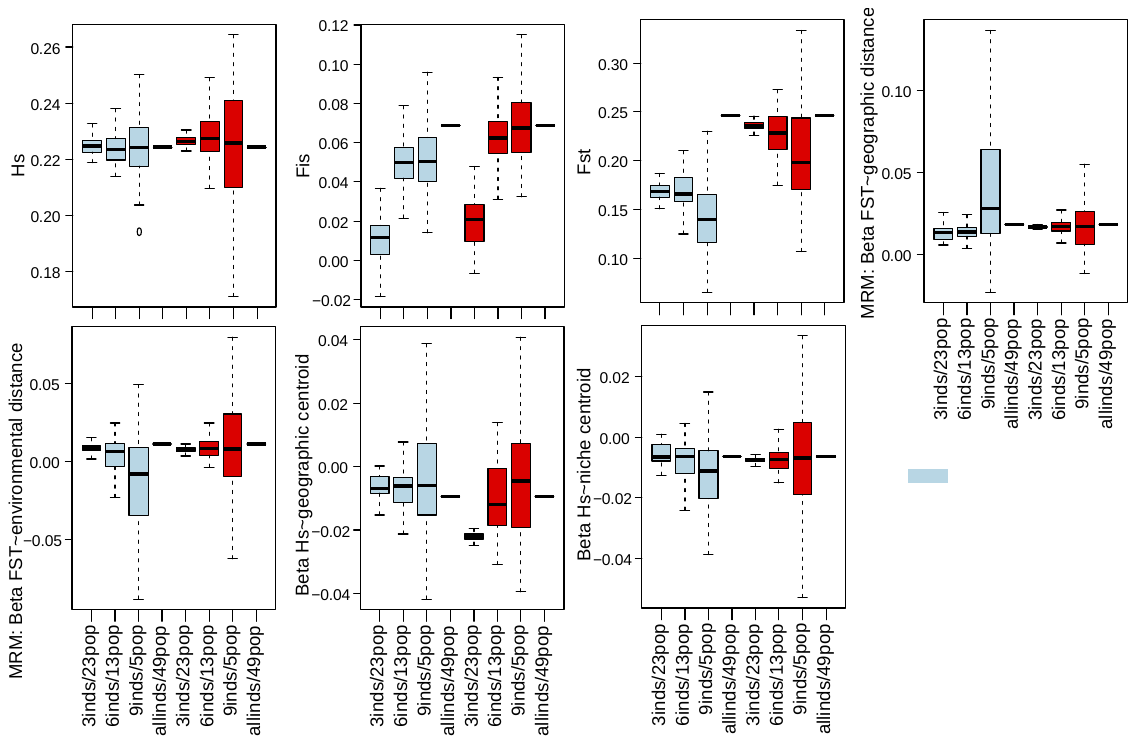

**FIG S5** Trade off between the number of individuals and the number of populations sampled for all summary statistics using the 50K dataset for *Zea mays* ssp *parviglumis* and *Zea mays* ssp *mexicana* populations. We tested the effect of sampling more individuals in few populations and few individuals in many populations.

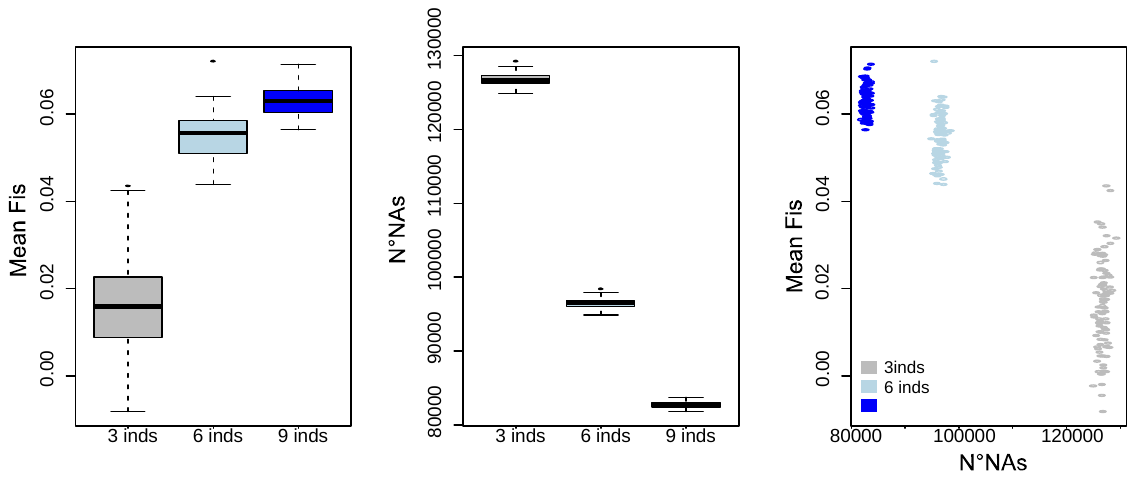

**FIG S6.** Estimations of mean F_IS_ across populations with different sampling size (3, 6 and 9 random individuals sampled per population). The left plot shows the distribution across 100 replicates. The Second plot shows the number of NAs across each sample. The third plot shows the correlation between the number of NAs and the estimation of F_IS_.

**Supporting tables**

**All supporting tables are found in the excel document supporting_tables_1to6.xlsx. Each table is found in a different sheet.**

**Table S1.** Mean, minimum and maximum values estimated for summary statistics across 1,000 replicates of different sampling designs. The real estimates are found in Table 1.

**Table S2.** Tukey tests showing the sampling designs that were significantly different (p-values are rounded, so 0 are values o p<0.00001).

**Table S3.** Number of significant differences between sampling designs. Barplots showing the counts can be found in Figure 4.

**Table S4.** Tests showing a significant deviation respect the real dataset.

**Table S5.** Percentage of false positive associations obtained for different sampling design replicates. False positives were considered when the β sign was opposit to the one estimated for the “real” dataset.

**Table S6.** Number of shared outlier SNPs between replicates and the real dataset.

**Custom R scripts that allow subsampling the datsets. For any inquiries please contact**

library(adegenet)

library(hierfstat)

library(ecodist)

library(BEDASSLE)

library(gplots)

num_inds <- function(N=3, replicates=10, especie="parviglumis",base="genomic", dataset="50K_genind.R"){

load(dataset)

d_genind <- dataset$genind

d_hierfstat <- dataset$hierfstat

d.frame <- data.frame(inds=rownames(d_genind$tab),

pop=d_genind$pop,num=1:nrow(d_genind$tab))

#populations <- as.character(unique(d.frame$pop))

if(especie=="mexicana"){

populations <- rownames(d_genind$other)[which(d_genind$other$Subespecie=="mexicana")]

}

if(especie=="parviglumis"){

populations <- rownames(d_genind$other)[which(d_genind$other$Subespecie=="parviglumis")]

}

if(especie=="all"){

populations <- rownames(d_genind$other)

}

locations <- d_genind$other

locations <- locations[populations,]

lon <-median(locations$lon)

lat <-median(locations$lat)

locations$geo_cent <- sqrt((locations$lon-lon)^2+(locations$lat-lat)^2)

hs_range <- NULL

fis_range <- NULL

fst_range <- NULL

hac <- NULL

hac_nic <- NULL

mantel_geo <- NULL

mantel_env <- NULL

MRM_env <- NULL

MRM_geo <- NULL

for(j in 1:replicates){

vec_inds <- NULL

for(i in 1:length(populations)){

pop <- populations[i]

inds <- d.frame[which(d.frame$pop==pop),]

inds <- inds$num[sample(1:nrow(inds),size = N,replace = F)]

vec_inds <- c(vec_inds,inds)

}

temp_genind <- d_genind[vec_inds,]

#temp_hierstat <- genind2hierfstat(temp_genind)

temp_hierstat <- d_hierfstat[vec_inds,]

temp_hierstat$pop <- factor(as.character(temp_hierstat$pop))

sumaries <- basic.stats(temp_hierstat)

hs_temp <- colMeans(sumaries$Hs,na.rm = T)

hs_range <- c(hs_range,as.vector(mean(hs_temp,na.rm = T)))

plot(locations$geo_cent,hs_temp,

xlab="geographic centroid",

ylab="Hs",main=paste("rep",j,"individuals",N))

fit <- lm(hs_temp~locations$geo_cent)

abline(fit)

fit <- summary(fit)

hac <- c(hac,fit$coefficients[2,1])

plot(locations$niche_centroid,hs_temp,

xlab="Niche centroid",ylab="Hs",main=paste("rep",j,"individuos",N))

fit <- lm(hs_temp~locations$niche_centroid)

abline(fit)

fit <- summary(fit)

hac_nic <- c(hac_nic,fit$coefficients[2,1])

fis_temp <- colMeans(sumaries$Fis,na.rm = T)

fis_range <- c(fis_range,as.vector(mean(fis_temp,na.rm = T)))

if(base=="microsatellite"){

fst <- pairwise.fst(temp_genind)

} else{

temp_genepop <- genind2genpop(temp_genind)

alleleA <- temp_genepop$tab[,seq(1,ncol(temp_genepop$tab)-1,2)]

alleleB <- temp_genepop$tab[,seq(2,ncol(temp_genepop$tab),2)]

counts <- alleleA+alleleB

fst <- as.dist(calculate.all.pairwise.Fst(alleleA,counts))

}

fst_range <- c(fst_range,as.vector(sumaries$overall["Fst"]))

geo <- dist(locations[c("lon","lat")])

env <- dist(locations$PC1)

plot(geo,fst,ylab="Fst",xlab="Geographic distance",main=paste("rep",j,"individuals",N))

r <- mantel(formula = fst~geo,nperm = 1000)

r <- r[1]

mantel_geo <- c(mantel_geo,r)

plot(env,fst,ylab="Fst",xlab="PC1 distance",main=paste("rep",j,"individuals",N))

r <- mantel(formula = fst~env,nperm = 1000)

r <- r[1]

mantel_env <- c(mantel_env,r)

mrm <- MRM(formula = fst~env+geo,nperm = 1000)

mrmenv <- mrm$coef["env","fst"]

mrmgeo <- mrm$coef["geo","fst"]

MRM_env <- c(MRM_env,mrmenv)

MRM_geo <- c(MRM_geo,mrmgeo)

print(paste("sp=",especie," inds N=",N,":",j))

if(j==333 | j==666){

sumaries <- list(hs_range=hs_range,

fis_range=fis_range,

fst_range=fst_range,

mantel_geo=mantel_geo,

mantel_env=mantel_env,

MRM_geo=MRM_geo,

MRM_env=MRM_env,

hac=hac,hac_nic=hac_nic)

save(sumaries,file = "temp_inds.R")

}

}

sumaries <- list(hs_range=hs_range,

fis_range=fis_range,

fst_range=fst_range,

mantel_geo=mantel_geo,

mantel_env=mantel_env,

MRM_geo=MRM_geo,

MRM_env=MRM_env,

hac=hac,hac_nic=hac_nic)

return(sumaries)

}

num_loc <- function(N=100, replicates=1000, especie="all",base="genomic", dataset="50K_genind.R"){

load(dataset)

d_genind <- dataset$genind

d_hierfstat <- dataset$hierfstat

d.frame <- data.frame(inds=rownames(d_genind$tab),

pop=d_genind$pop,num=1:nrow(d_genind$tab))

populations <- as.character(unique(d.frame$pop))

if(especie=="mexicana"){

populations <- rownames(d_genind$other)[which(d_genind$other$Subespecie=="mexicana")]

}

if(especie=="parviglumis"){

populations <- rownames(d_genind$other)[which(d_genind$other$Subespecie=="parviglumis")]

}

if(especie=="all"){

populations <- rownames(d_genind$other)

}

locations <- d_genind$other

locations <- locations[populations,]

lon <-median(locations$lon)

lat <-median(locations$lat)

locations$geo_cent <- sqrt((locations$lon-lon)^2+(locations$lat-lat)^2)

hs_range <- NULL

fis_range <- NULL

fst_range <- NULL

hac <- NULL

hac_nic <- NULL

mantel_geo <- NULL

mantel_env <- NULL

MRM_env <- NULL

MRM_geo <- NULL

j <- 1

while(j <= replicates){

print(j)

vec_loc <- NULL

vec_loc_hierf <- NULL

loci <- names(d_genind$loc.n.all)

loci <- sample(loci,size = N,replace = F)

ja <- NULL

for(i in 1:length(loci)){

temp_loc <- paste(loci[i],".",sep = "")

temp_loc <- grep(pattern = temp_loc,x = colnames(d_genind$tab),fixed = T)

ja <- c(ja, length(temp_loc))

vec_loc <- c(vec_loc,temp_loc)

}

vec_loc <- vec_loc[order(vec_loc)]

temp_genind <- d_genind[,vec_loc]

vec_pop <- NULL

for(i in 1:length(populations)){

#print(i)

#print(populations[i])

vec_pop <- c(vec_pop,grep(paste(populations[i],"_",sep = ""),rownames(temp_genind$tab),fixed = T))

\# print(rownames(temp_genind$tab)[grep(paste(populations[i],"_",sep = ""),rownames(temp_genind$tab),fixed = T)])

\# print("")

}

dup <- which(duplicated(vec_pop))

if(length(dup)>0){

vec_pop<- vec_pop[-dup]

}

temp_genind <- temp_genind[vec_pop,]

alleles.name <- toupper(names(table(unlist(temp_genind@all.names))))

if(length(alleles.name)!=4){next}

#temp_hierstat <- genind2hierfstat(temp_genind)

loc_hier <- c("pop",as.vector(names(temp_genind$all.names)))

temp_hierstat <- d_hierfstat[vec_pop,loc_hier]

temp_hierstat$pop <- factor(temp_hierstat$pop)

summaries <- basic.stats(temp_hierstat)

hs_temp <- colMeans(summaries$Hs,na.rm = T)

hs_range <- c(hs_range,mean(hs_temp,na.rm = T))

locations <- locations[names(hs_temp),]

plot(locations$geo_cent,hs_temp,

xlab="geographic centroid",

ylab="Hs",main=paste("rep",j,"loci",N))

fit <- lm(hs_temp~locations$geo_cent)

abline(fit)

fit <- summary(fit)

hac <- c(hac,fit$coefficients[2,1])

plot(locations$niche_centroid,hs_temp,

xlab="Niche centroid",ylab="Hs",main=paste("rep",j,"loci",N))

fit <- lm(hs_temp~locations$niche_centroid)

abline(fit)

fit <- summary(fit)

hac_nic <- c(hac_nic,fit$coefficients[2,1])

fis_temp <- colMeans(summaries$Fis,na.rm = T)

fis_range <- c(fis_range,mean(hs_temp,na.rm = T))

if(base=="microsatellite"){

fst <- pairwise.fst(temp_genind)

} else{

temp_genepop <- genind2genpop(temp_genind)

alleleA <- temp_genepop$tab[,seq(1,ncol(temp_genepop$tab)-1,2)]

alleleB <- temp_genepop$tab[,seq(2,ncol(temp_genepop$tab),2)]

counts <- alleleA+alleleB

fst <- as.dist(calculate.all.pairwise.Fst(alleleA,counts))

}

fst_range <- c(fst_range,summaries$overall["Fst"])

geo <- dist(locations[c("lon","lat")])

env <- dist(locations$PC1)

plot(geo,fst,ylab="Fst",xlab="Geographic distance",main=paste("rep",j,"loci",N))

r <- mantel(formula = fst~geo,nperm = 1000)

r <- r[1]

mantel_geo <- c(mantel_geo,r)

plot(env,fst,ylab="Fst",xlab="PC1 distance",main=paste("rep",j,"loci",N))

r <- mantel(formula = fst~env,nperm = 1000)

r <- r[1]

mantel_env <- c(mantel_env,r)

mrm <- MRM(formula = fst~env+geo,nperm = 1000)

mrmenv <- mrm$coef["env","fst"]

mrmgeo <- mrm$coef["geo","fst"]

MRM_env <- c(MRM_env,mrmenv)

MRM_geo <- c(MRM_geo,mrmgeo)

print(paste("sp=",especie," loc N=",N,":",j))

j <- j+1

if(j==333 | j==666){

sumaries <- list(hs_range=hs_range,

fis_range=fis_range,

fst_range=fst_range,

mantel_geo=mantel_geo,

mantel_env=mantel_env,

MRM_geo=MRM_geo,

MRM_env=MRM_env,

hac=hac,hac_nic=hac_nic)

save(sumaries,file = "temp_loc.R")

}

}

sumaries <- list(hs_range=hs_range,

fis_range=fis_range,

fst_range=fst_range,

mantel_geo=mantel_geo,

mantel_env=mantel_env,

MRM_geo=MRM_geo,

MRM_env=MRM_env,

hac=hac,hac_nic=hac_nic)

return(sumaries)

}

num_pops <- function(N=5, replicates=1000, especie= "all" ,base="genomic", dataset="50K_genind.R"){

load(dataset)

d_genind <- dataset$genind

d_hierfstat <- dataset$hierfstat

d.frame <- data.frame(inds=rownames(d_genind$tab),

pop=d_genind$pop,num=1:nrow(d_genind$tab))

populations <- as.character(unique(d.frame$pop))

if(especie=="mexicana"){

populations <- rownames(d_genind$other)[which(d_genind$other$Subespecie=="mexicana")]

}

if(especie=="parviglumis"){

populations <- rownames(d_genind$other)[which(d_genind$other$Subespecie=="parviglumis")]

}

if(especie=="all"){

populations <- rownames(d_genind$other)

}

locations <- d_genind$other

locations <- locations[populations,]

lon <-median(locations$lon)

lat <-median(locations$lat)

locations$geo_cent <- sqrt((locations$lon-lon)^2+(locations$lat-lat)^2)

hs_range <- NULL

fis_range <- NULL

fst_range <- NULL

hac <- NULL

hac_nic <- NULL

mantel_geo <- NULL

mantel_env <- NULL

MRM_env <- NULL

MRM_geo <- NULL

for(j in 1:replicates){

temp_populations <- sample(populations,size = N,replace = F)

vec_pops <- NULL

vec_loc <- NULL # here loc means locations not loci

for(i in 1:length(temp_populations)){

pop <- temp_populations[i]

pops <- d.frame[which(d.frame$pop==pop),]

pops <- pops$num

vec_pops <- c(vec_pops,pops)

vec_loc <- c(vec_loc,which(rownames(locations)==pop))

}

temp_genind <- d_genind[vec_pops,]

#temp_hierstat <- genind2hierfstat(temp_genind)

temp_hierstat <- d_hierfstat[vec_pops,]

temp_hierstat$pop <- factor(temp_hierstat$pop)

sumaries <- basic.stats(temp_hierstat)

hs_temp <- colMeans(sumaries$Hs,na.rm = T)

hs_range <- c(hs_range,mean(hs_temp,na.rm = T))

temp_locations <- locations[vec_loc,]

temp_locations <- data.frame(temp_locations)

temp_locations <- temp_locations[names(hs_temp),]

plot(temp_locations$geo_cent,hs_temp,

xlab="geographic centroid",

ylab="Hs",main=paste("rep",j,"individuals",N))

fit <- lm(hs_temp~temp_locations$geo_cent)

abline(fit)

fit <- summary(fit)

hac <- c(hac,fit$coefficients[2,1])

plot(temp_locations$niche_centroid,hs_temp,

xlab="Niche centroid",ylab="Hs",main=paste("rep",j,"pop",N))

fit <- lm(hs_temp~temp_locations$niche_centroid)

abline(fit)

fit <- summary(fit)

hac_nic <- c(hac_nic,fit$coefficients[2,1])

fis_temp <- colMeans(sumaries$Fis,na.rm = T)

fis_range <- c(fis_range,mean(fis_temp,na.rm = T))

if(base=="microsatellite"){

fst <- pairwise.fst(temp_genind)

} else{

temp_genepop <- genind2genpop(temp_genind)

alleleA <- temp_genepop$tab[,seq(1,ncol(temp_genepop$tab)-1,2)]

alleleB <- temp_genepop$tab[,seq(2,ncol(temp_genepop$tab),2)]

counts <- alleleA+alleleB

fst <- as.dist(calculate.all.pairwise.Fst(alleleA,counts))

}

fst_range <- c(fst_range,summaries$overall["Fst"])

geo <- dist(temp_locations[c("lon","lat")])

env <- dist(temp_locations$PC1)

plot(geo,fst,ylab="Fst",xlab="Geographic distance",main=paste("rep",j,"pop",N))

r <- mantel(formula = fst~geo,nperm = 1000)

r <- r[1]

mantel_geo <- c(mantel_geo,r)

plot(env,fst,ylab="Fst",xlab="PC1 distance",main=paste("rep",j,"pop",N))

r <- mantel(formula = fst~env,nperm = 1000)

r <- r[1]

mantel_env <- c(mantel_env,r)

mrm <- MRM(formula = fst~env+geo,nperm = 1000)

mrmenv <- mrm$coef["env","fst"]

mrmgeo <- mrm$coef["geo","fst"]

MRM_env <- c(MRM_env,mrmenv)

MRM_geo <- c(MRM_geo,mrmgeo)

print(paste("sp=",especie," pops N=",N,":",j))

if(j==333 | j==666){

sumaries <- list(hs_range=hs_range,

fis_range=fis_range,

fst_range=fst_range,

mantel_geo=mantel_geo,

mantel_env=mantel_env,

MRM_geo=MRM_geo,

MRM_env=MRM_env,

hac=hac,hac_nic=hac_nic)

save(sumaries,file = "temp_pops.R")

}

}

sumaries <- list(hs_range=hs_range,

fis_range=fis_range,

fst_range=fst_range,

mantel_geo=mantel_geo,

mantel_env=mantel_env,

MRM_geo=MRM_geo,

MRM_env=MRM_env,

hac=hac,hac_nic=hac_nic)

return(sumaries)

}

tradeoffs <- function(N=3, Npop=10,replicates=1000,especie="all", base="genomic", dataset="50K_genind.R"){

load(dataset)

d_genind <- dataset$genind

d_hierfstat <- dataset$hierfstat

d.frame <- data.frame(inds=rownames(d_genind$tab),

pop=d_genind$pop,num=1:nrow(d_genind$tab))

populations <- as.character(unique(d.frame$pop))

if(especie=="mexicana"){

populations <- rownames(d_genind$other)[which(d_genind$other$Subespecie=="mexicana")]

}

if(especie=="parviglumis"){

populations <- rownames(d_genind$other)[which(d_genind$other$Subespecie=="parviglumis")]

}

if(especie=="all"){

populations <- rownames(d_genind$other)

}

populations <- populations[sample(1:length(populations),size = Npop,replace = F)]

locations <- d_genind$other

locations <- locations[populations,]

lon <-median(locations$lon)

lat <-median(locations$lat)

locations$geo_cent <- sqrt((locations$lon-lon)^2+(locations$lat-lat)^2)

hs_range <- NULL

fis_range <- NULL

fst_range <- NULL

hac <- NULL

hac_nic <- NULL

mantel_geo <- NULL

mantel_env <- NULL

MRM_env <- NULL

MRM_geo <- NULL

for(j in 1:replicates){

vec_inds <- NULL

for(i in 1:length(populations)){

pop <- populations[i]

inds <- d.frame[which(d.frame$pop==pop),]

inds <- inds$num[sample(1:nrow(inds),size = N,replace = F)]

vec_inds <- c(vec_inds,inds)

}

temp_genind <- d_genind[vec_inds,]

#temp_hierstat <- genind2hierfstat(temp_genind)

temp_hierstat <- d_hierfstat[vec_inds,]

temp_hierstat$pop <- factor(as.character(temp_hierstat$pop))

sumaries <- basic.stats(temp_hierstat)

hs_temp <- colMeans(sumaries$Hs,na.rm = T)

hs_range <- c(hs_range,mean(hs_temp,na.rm = T))

locations <- locations[names(hs_temp),]

plot(locations$geo_cent,hs_temp,

xlab="geographic centroid",

ylab="Hs",main=paste("rep",j,"individuals",N))

fit <- lm(hs_temp~locations$geo_cent)

abline(fit)

fit <- summary(fit)

hac <- c(hac,fit$coefficients[2,1])

plot(locations$niche_centroid,hs_temp,

xlab="Niche centroid",ylab="Hs",main=paste("rep",j,"individuos",N))

fit <- lm(hs_temp~locations$niche_centroid)

abline(fit)

fit <- summary(fit)

hac_nic <- c(hac_nic,fit$coefficients[2,1])

fis_temp <- colMeans(sumaries$Fis,na.rm = T)

fis_range <- c(fis_range,mean(fis_temp,na.rm = T))

if(base=="microsatellite"){

fst <- pairwise.fst(temp_genind)

} else{

temp_genepop <- genind2genpop(temp_genind)

alleleA <- temp_genepop$tab[,seq(1,ncol(temp_genepop$tab)-1,2)]

alleleB <- temp_genepop$tab[,seq(2,ncol(temp_genepop$tab),2)]

counts <- alleleA+alleleB

fst <- calculate.all.pairwise.Fst(alleleA,counts)

fst <- as.dist(fst)

}

fst_range <- c(fst_range,summaries$overall["Fst"])

geo <- dist(locations[c("lon","lat")])

env <- dist(locations$PC1)

plot(geo,fst,ylab="Fst",xlab="Geographic distance",main=paste("rep",j,"individuals",N))

r <- mantel(formula = fst~geo,nperm = 1000)

r <- r[1]

mantel_geo <- c(mantel_geo,r)

plot(env,fst,ylab="Fst",xlab="PC1 distance",main=paste("rep",j,"individuals",N))

r <- mantel(formula = fst~env,nperm = 1000)

r <- r[1]

mantel_env <- c(mantel_env,r)

mrm <- MRM(formula = fst~env+geo,nperm = 1000)

mrmenv <- mrm$coef["env","fst"]

mrmgeo <- mrm$coef["geo","fst"]

MRM_env <- c(MRM_env,mrmenv)

MRM_geo <- c(MRM_geo,mrmgeo)

print(paste("sp=",especie," trade N=",N,":",j))

if(j==333 | j==666){

sumaries <- list(hs_range=hs_range,

fis_range=fis_range,

fst_range=fst_range,

mantel_geo=mantel_geo,

mantel_env=mantel_env,

MRM_geo=MRM_geo,

MRM_env=MRM_env,

hac=hac,hac_nic=hac_nic)

save(sumaries,file = "temp_trade_offs.R")

}

}

sumaries <- list(hs_range=hs_range,

fis_range=fis_range,

fst_range=fst_range,

mantel_geo=mantel_geo,

mantel_env=mantel_env,

MRM_geo=MRM_geo,

MRM_env=MRM_env,

hac=hac,hac_nic=hac_nic)

return(sumaries)

}
